# Mutations in *PSEN1* predispose inflammation in an astrocyte model of familial Alzheimer’s disease through disrupted regulated intramembrane proteolysis

**DOI:** 10.1101/2024.09.09.611621

**Authors:** Oliver J. Ziff, Sarah Jolly, Jackie M. Casey, Lucy Granat, Satinder Samra, Nuria Seto-Salvia, Argyro Alatza, Leela Phadke, Benjamin Galet, Philippe Ravassard, Marie-Claude Potier, Nick C. Fox, John Hardy, Dervis Salih, Paul Whiting, Fiona Ducotterd, Rickie Patani, Selina Wray, Charles Arber

**Affiliations:** Department of Neuromuscular Diseases, UCL Queen Square Institute of Neurology, University College London, London, UK; Alzheimer’s Research UK UCL Drug Discovery Institute, University College London, London, UK; Department of Neurodegenerative Disease, UCL Queen Square Institute of Neurology, London, UK; Department of Clinical and Movement Neuroscience, Reta Lila Weston Institute, UCL Queen Square Institute of Neurology, London, UK; Sorbonne Université, Institut du Cerveau - Paris Brain Institute - ICM, Inserm, CNRS, APHP, Hôpital de la Pitié Salpêtrière, Paris, France; UK Dementia Research Institute at UCL, London, UK; Dementia Research Centre, UCL Queen Square Institute of Neurology, London, UK; Human Stem Cells and Neurodegeneration Laboratory, The Francis Crick Institute, London, UK

**Keywords:** iPSC, astrocyte, PSEN1, Alzheimer’s disease, inflammation, regulated intramembrane proteolysis

## Abstract

Mutations in *PSEN1* cause familial Alzheimer’s disease with almost complete penetrance. Age at onset is highly variable between different *PSEN1* mutations and even within families with the same mutation. Current research into late onset Alzheimer’s disease implicates inflammation in both disease onset and progression. PSEN1 is the catalytic subunit of γ-secretase, responsible for regulated intramembrane proteolysis of numerous substrates that include cytokine receptors. For this reason, we tested the hypothesis that mutations in *PSEN1* impact inflammatory responses in astrocytes, thereby contributing to disease progression.

Here, using iPSC-astrocytes, we show that PSEN1 is upregulated in response to inflammatory stimuli, and this upregulation is disrupted by pathological *PSEN1* mutations. Using transcriptomic analyses, we demonstrate that *PSEN1* mutant astrocytes have an augmented inflammatory profile in their basal state, concomitant with an upregulation of genes coding for regulated intramembrane proteolytic and robust activation of JAK-STAT signalling. Using JAK-STAT2 as an example signalling pathway, we show altered phosphorylation cascades in *PSEN1* mutant astrocytes, reinforcing the notion of altered cytokine signalling cascades. Finally, we use small molecule modulators of γ-secretase to confirm a role for PSEN1/γ-secretase in regulating the astrocytic response to inflammatory stimuli.

Together, these data suggest that mutations in *PSEN1* enhance cytokine signalling via impaired regulated intramembrane proteolysis, thereby predisposing astrocytic inflammatory profiles. These findings support a two-hit contribution of *PSEN1* mutations to fAD pathogenesis, not only impacting APP and Aβ processing but also altering the cellular response to inflammation.

## Introduction

Together with amyloid plaques and tau tangles, neuroinflammation represents one of the key pathological hallmarks of Alzheimer’s disease (AD). Both astrocytes and microglia proliferate and cluster around amyloid plaques [57], and alter their inflammation-associated transcriptomic signatures [23, 29]. It remains unclear whether inflammation is protective or harmful, however, evidence suggests that the innate immune response may link the long preclinical amyloid phase of AD to the clinical tau phase of AD [33, 36]. Recent data support a central role for astrocyte dysfunction early in AD pathogenesis, for example, via an association of astrocyte-enriched SNPs with early amyloid pathology [67], as well as changes observed to both MOA-B PET tracer positivity [16] and plasma levels of GFAP [44] prior to amyloid plaque formation. The astrocyte response to inflammatory factors released by microglia has been shown to have a central role in mediating neurodegeneration [37], although, it has been suggested that astrocyte responses may play a crucial protective role in AD [41].

Familial Alzheimer’s disease (fAD) is a rare, inherited, autosomal dominant form of AD caused by mutations in *PSEN1*, *PSEN2* and *APP* [18, 35, 60]. *PSEN1/2* encode the catalytic subunit of γ-secretase, which cleaves APP to generate Aβ, the major component of amyloid plaques [25]. Mutations in *PSEN1* destabilise the enzyme-substrate complex, thereby releasing longer forms of Aβ prior to complete processing [48, 56, 63]. These longer forms, such as Aβ42 and Aβ43, are aggregation prone and are found in plaque cores. One hypothesis linking fAD and late onset AD (LOAD) states that fAD is associated with an increased production of plaque-forming Aβ peptides, whereas LOAD is associated with decreased clearance of Aβ; this is supported by kinetic *in vivo* data [42, 49].

Evidence from human iPSC-derived neuronal models (which can be considered foetal-like in their maturation status) and young mouse knock-in models suggest that mutations in *PSEN1* lead to a constitutive alteration in Aβ production, i.e. from birth [4, 61]. Despite this observation, the clinical onset of fAD affects individuals in their third to seventh decades of life [51, 52], and shows considerable heterogeneity in disease onset, progression and clinical presentation, even within families with the same mutation. This suggests that, even in fAD, additional genetic or environmental factors contribute to disease onset and progression [51].

In addition to APP, γ-secretase cleaves more than 150 known substrates [19, 22], including the key inflammatory cytokine receptors IL1R [14] and TNFR1 [11]. Cell signalling can be controlled by γ-secretase-mediated cleavage, such that active signalling stubs of receptors can be cleaved for degradation (cessation of signal) or translocation (potentiation of signal) [43]. Thereby, it has been suggested that γ-secretase-mediated cleavage can modulate the inflammatory response. Examples of this include the cleavage of LRP1 and SIRPα that have been shown reduce or enhance inflammation respectively [38, 68]. More recently, a study of γ-secretase inhibition/deficiency in iPSC-derived microglia and model mice, elegantly demonstrated that γ-secretase regulates key microglial genes and orchestrates cell state transitions *in vivo* [26]. In contrast, little is known about the effect of fAD mutations in *PSEN1* on cytokine responses and cell state transitions.

Here, we hypothesised that *PSEN1* mutations not only increase the production of aggregation-prone species of Aβ, but also alter the cellular responses to inflammatory cytokines through impaired cytokine receptor cleavage. We therefore established a patient-derived iPSC-astrocyte model to investigate the effect of *PSEN1* mutations on glial responses to inflammatory stimuli.

## Methods

### Cell culture and treatments

All iPSC lines used in this study have been characterised and described previously. This is with the exception of the isogenic corrected R278 line following the protocol described previously [34] using the CRISPR/Cas9 technology [10]. Briefly, iPSC cultured in mTeSR-Plus medium were dissociated as single cells with Accutase (Stem Cell Technologies). One million cells were then transfected (4D Nucleofector system, Core Unit AAF-1002B and X unit AAF-1002; Lonza, Switzerland) with a ribonucleoprotein (RNP) complex and 500 pmol of ssODN (single stranded oligodeoxynucleotide) matrix for homologous recombination (TCTCAAATACTTACAGGAGTAAATGAGAGCTGGAAAAAGCGTTTCATTTCTCTCCTGTGCAGTTTCAACCAGCATACGAAGTGGACCTTTCGGACACAAA). The RNP complex was prepared with equimolar concentrations of crRNA (TATGCTGGTTGAAACAGCTC) and tracrRNA-ATT0550 (225 pmol of each RNA), with 120 pmol of Cas9 protein (Integrated DNA Technologies (IDT), Iowa, USA). After transfection, all cells were plated and the day after ATT0550-positive cells were sorted by FACS (MoFlo Astrios, Beckman Coulter; ICV-Cytometrie, ICM, Paris, France) and plated at low concentrations on petri dishes coated with Laminin521 (Stem Cell Technologies) in mTeSR-Plus medium supplemented with Clone R (Stem Cell Technologies). Seven to 10 days later, individual iPSC clones were picked and expanded in 96-well plates. Each clone was duplicated for either DNA analysis or freezing. Between 100 and 200 clones were picked up. Genetically engineered clones were screened after DNA extraction by PCR amplification of the targeted genomic region and sequencing using the following primers: FWD TCCTCCCTACCACCCATTTAC and REV GGAGTTCCAGGAATGCTGTG (Fig. S1). Corrected clones were then amplified and characterized at the molecular level. Potential off-target sites with one, two or three mismatches were screened by Sanger sequencing (Eurofins Genomics). No off-target mutations were found in the selected clones. Corrected clones were tested for recurrent genomic abnormalities by the ICS-digital™ PSC test (Stem Genomics, Montpellier, France).

All details of iPSC lines are described in the table below. iPSCs were cultured in Essential 8 media on Geltrex substrate and passaged using EDTA mechanical passaging. All components were purchased from ThermoFisher and all recombinant proteins were Peprotech, unless specified. Sanger sequencing was performed with SourceBio. Low coverage whole genome sequencing was performed in collaboration with UCL Genomics. Reads were divided into 1000kb bins and the QDNASeq package was employed that included smoothing and control for mapability [54].

Human foetal astrocytes were purchased ScienCell Research Laboratories (catalogue #1800) and maintained and passaged in an identical manner to the iPSC-derived astrocytes.

To generate iPSC-derived cortical neurons, we followed established protocols [1, 3–5, 66]. In brief, neural induction was performed using SB431542 (10μM, Tocris) and dorsomorphin (1μM, Tocris) in N2B27 media. N2B27 media is composed of 50% DMEM-F12, 50% Neurobasal with the addition of 0.5X N2 supplement, 0.5X B27 supplement, 0.5X L-glutamine, 0.5X NEAA, 0.5X pen/strep (25U), 1:1000 β-mercaptoethanol, and insulin at 25U. 100 DIV was used as the final time-point.

For the generation of iPSC-astrocytes we followed established protocols [2, 24, 58]. Glial precursor cells (GPCs) were enriched from neuronal cultures at 80 DIV by regular passaging using EDTA and the addition of FGF2 (10ng/ml, Peprotech) to the N2B27 media. A gliogenic switch occurred around 110 DIV. To generate mature astrocytes, GPCs of at least 150 DIV were subjected to two weeks maturation with 10ng/ml BMP4 (Peprotech) and 10ng/ml LIF (Sigma) in N2B27 media.

iPSC-microglia were generated using established protocols [2, 17]. Briefly, haematopoiesis was induced in iPSC-derived embryoid bodies via treatment with 50ng/ml BMP4, 50ng/ml VEGF and 20ng/ml SCF. Myeloid patterning was performed using MCSF (100ng/ml) and IL3 (25ng/ml) in XVIVO media (Lonza). Microglia progenitors were harvested from the embryoid bodies supernatants and matured in media containing MCSF (25ng/ml), IL34 (100ng/ml) and TGFβ (5ng/ml), with a final two-day treatment in CX3CL1 (100ng/ml) and CD200 (100ng/ml).

Conditioned media was collected after 48 hours and spun at 2,000G to remove cell debris, prior to using for Western blotting, ELISA, cytokine arrays or LDH assays.

Cell treatments included TIC, where TNFα (33ng/ml, Peprotech), IL1α (3ng/ml Peprotech) and C1Q (400ng/ml, Merck) were used in combination for 24 hours. γ-secretase modulation was performed with E2012 (GSM, 10μM, MedChemExpress). γ-secretase inhibition was performed using DAPT (GSI, 10μM, Tocris). For treatments with GSI/GSM in combination with TIC a 1-hour pre-treatment with GSI/GSM was performed prior to the addition of TIC.

### Immunocytochemistry and high content imaging

Cells were fixed using 4% paraformaldehyde for 15 minutes. Three washes were performed in PBS with 0.3% triton-x-100 (PBSTx). Blocking was done for 20 minutes using 3% bovine serum albumin in PBSTx. Primary antibodies (Table 2) were incubated in blocking solution overnight at 4°C. Secondary antibodies (AlexaFluor) were incubated for 1 hour in blocking solution after 3 washes in PBSTx. DAPI was used as a counterstain with final washes. Imaging was done using the high content imaging system Opera Phenix (Perkin Elmer) and analysis was done on the Columbus software. No post-hoc image processing was performed.

### Western Blotting

Cells were lysed in RIPA buffer containing phosphatase and protease inhibitors (Roche) and spun to remove insoluble debris. Protein content was quantified using the BioRad BCA assay. Samples were denatured in LDS with DTT and loaded onto 4-12% precast polyacrylamine gels. Gels were transferred to nitrocellulose membranes and then blocked in PBS with 0.1% Tween20 (PBSTw) with 3% bovine serum albumin. Primary antibodies (Table 2) were incubated in blocking solution overnight, before 3 washes in PBSTw and secondary antibody incubation (Licor) for 1 hour. Membranes were imaged using a Licor Odyssey fluorescent imaging system. Quantification was done on the ImageStudio software.

### Quantitative PCR

For qPCR, cells were lysed in Trizol and RNA was isolated following the manufacturer’s protocols. cDNA was synthesised from 1-2μg of RNA using Superscript IV with random hexamer primers and RNAse OUT. qPCR was run using Power SYBR green on an Agilent Aria MX machine with annealing at 60°C. Samples were analysed relative to the housekeeping gene *RPL18a* and relative expression was quantified using the ΔΔCT method. qPCR primers are listed in Table 3.

### RNA sequencing

Cells were lysed in Trizol and RNA was harvested using the Monarch total RNA miniprep kit (NEB). Poly(A)+ selected sense-stranded RNA sequencing libraries were prepared using the KAPA mRNA Hyper Prep (Roche), with 500 ng of total RNA as input. Libraries were sequenced on the NextSeq 2000 platform for neuronal data and NovaSeq 6000 for astrocyte data. A mean of 189 (range 97–284) million 55 bp paired-end strand-specific reads were sequenced per sample. mRNA sequencing reads were processed using the nf-core/rna-seq v3.11.2 pipeline [15]. Raw reads underwent adaptor trimming with Trim Galore, removal of ribosomal RNA with SortMeRNA, alignment to Ensembl GRCh38.99 human reference genome using splice-aware aligner, STAR v2.6.1 and BAM level quantification with Salmon to enable transcript level counting. Detailed quality control of raw and aligned reads were assessed by utilizing FastQC, RSeQC, Qualimap, dupRadar, Preseq and MultiQC tools. All libraries generated in this study had <0.1% rRNA, <0.3% mismatch error, and >85% strandedness.

### Transcriptomic analysis

STAR aligned and Salmon quantified transcript abundance were summarised at the gene-level using tximport in R v4.2.0. Gene counts were normalized and transformed using the variance stabilizing transformation function in DESeq2[39]. These transformed values were utilized in the <u>principal component analysis</u> (PCA) and unsupervised hierarchical clustering. We examined the astrocyte transcriptomic identities of iPSC-astrocytes using the ComplexHeatmap package based on the expression of canonical neuronal and <u>glial cell</u> type markers.

Differential gene expression analysis was fitted using DESeq2 and normalised using the mean of ratios. Results of *PSEN1* mutant iPSC-astrocytes were generated by comparing the *PSEN1* mutant versus non mutant control samples using the Wald test. To examine the effect of TIC on gene expression we compared the TIC treated versus untreated samples. Results for *PSEN1* mutant astrocytes were correlated with TIC treated astrocytes by matching the Wald test statistic for each gene followed by Pearson correlation. In all analyses, genes were considered differentially expressed at FDR < 0.05. Significantly up- and down-regulated differentially expressed genes were used as input to functional over-representation analyses to identify enriched pathways using g:Profiler2. g:Profiler2 searches the following data sources: Gene Ontology (GO; molecular functions, biological processes and cellular components), KEGG, REAC, WikiPathways, CORUM and Human Phenotype Ontology. g:Profiler2 reports the hypergeometric test p-value with an adjustment for multiple testing using the Bonferroni correction. Over-represented function categories are plotted in bar charts, where the top significant terms were manually curated by removing redundant terms. The decoupleR package was used to estimate PROGENy signalling pathway activities and DoRothEA TF regulon activities inferred from gene expression changes[6]. PROGENy and DoRothEA weights are based on perturbation experiments that are not specific to motor neurons. Their signalling pathways may activate diverse downstream gene expression programmes depending on the cell type and perturbing agent utilised[55].

### Cytokine array

The Proteome Profiler Human XL Cytokine Array Kit (R&D Systems) was used, as per the manufacturer’s instructions. Conditioned media from three independent batches were pooled (160μl x 3) to generate an average reading per genotype and per condition.

### Lactate dehydrogenase assay

Lactate dehydrogenase assays were performed on conditioned media to quantify cell death. The Abcam LDH assay kit (ab65393) was used following the manufacturers instruction and the final absorbance was read on a Tecan SPARK 10M plate reader at 450nm.

### ELISAs

Aβ40 and Aβ42 were quantified using the Meso Scale Discovery V-Plex Aβ peptide panel (6E10). Samples were diluted 2-fold and quantification was performed on an MSD Sector 6000.

To quantify secreted TNFR1, we used the Quantikine Human TNF RI/TNFRSF1A ELISA from R&D Systems (DRT100), following the manufacturer’s protocols. Absorbance was measured on a Tecan SPARK 10M plate reader.

### Statistics

Data were collected and analysed using Microsoft Excel and GraphPad Prism. Normality was tested using D’Agostino and Pearson test. Comparisons between two groups (qPCR and Western blot data) were performed using two tailed t-tests. Data is shown as p<0.05 = *, p<0.01 = **, p<0.001 = *** and p<0.0001 = ****.

Outliers were identified if they had a z-score of above 3 or below −3. Z scores were calculated as (sample - mean)/standard deviation, for normally distributed data.

## Results

### Development of an iPSC-derived astrocyte model of fAD

To test the hypothesis that mutations in *PSEN1* alter glial inflammatory responses, we generated patient iPSC-derived astrocyte models of fAD following an established serum-free protocol [24, 58]. We focused our investigations on astrocytes, as these cells represent an immune competent cell type that expresses high levels of both APP and PSEN1 (Fig S2). We analysed three fAD patient-derived lines (with the int4del, Y115H and R278I mutations in *PSEN1*) with six healthy control lines (two of which were isogenic corrected lines – Table 1). As an established inflammatory cue, we use TNFα, IL1α and C1Q [37], hereafter referred to as TIC and performed deep RNA sequencing using poly(A) strand specific sequencing on paired samples (see Methods).

**Table 1.**
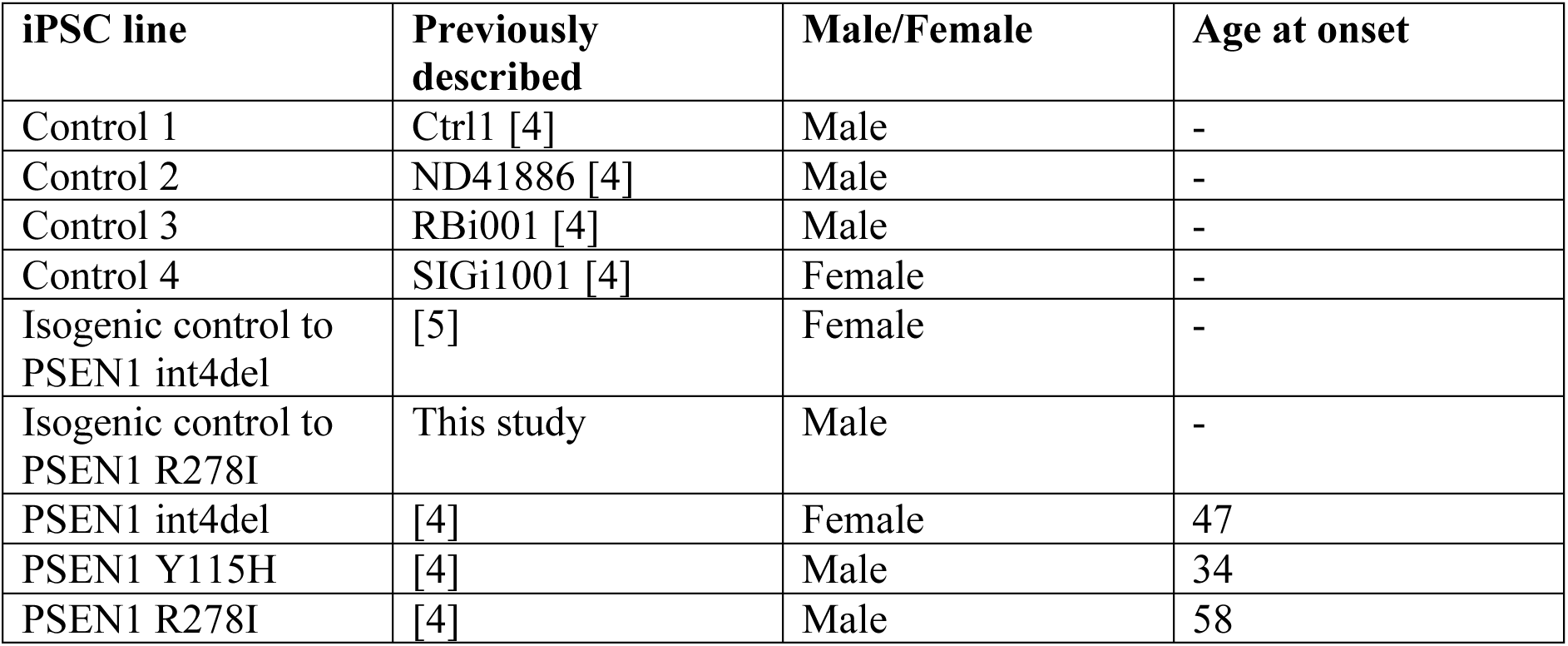
Cell lines used in this study.

**Table 2.** Antibodies for ICC and Western blotting.

| Antigen | Species | Company<br>(catalogue number) | RRID |
| --- | --- | --- | --- |
| SOX9 | rabbit | Abcam ab185966 | RRID:AB_2728660 |
| GFAP | mouse | Sigma G6171 | RRID:AB_1840893 |
| ALDH1L1 | rabbit | Abcam<br>ab87117 | RRID:AB_10712968 |
| Phalloidin | - | SantaCruz<br>Sc-363797 |  |
| pMAPK | rabbit | CST 9101 | RRID:AB_331646 |
| MAPK | rabbit | CST 9102 | RRID:AB_330744 |
| pSTAT2 | rabbit | CST 88410 | RRID:AB_2800123 |
| STAT2 | rabbit | Abcam Ab32367 | RRID:AB_778098 |
| pSTAT3 | rabbit | CST 9145 | RRID:AB_2491009 |
| STAT3 | rabbit | CST 4904 | RRID:AB_331269 |
| pNFκB | rabbit | CST 3033 | RRID:AB_331284 |
| NFκB | rabbit | CST 8242 | RRID:AB_10859369 |
| APP | rabbit | Thermo A8717 | RRID:AB_258409 |
| PSEN1 | rat | Millipore MAB1563 | RRID: B_11215630 |
| PSEN2 | rabbit | Cell Signalling<br>Technology 9979 | RRID:AB_10829910 |
| Actin | mouse | Sigma<br>A1978 | RRID:AB_476692 |
| GAPDH | mouse | Ambion AM4300 | RRID:AB_437392 |
| sAPPβ | rabbit | IBL JP18957 | RRID:AB_1630824 |
| sAPP total | mouse | (22c11) Millipore<br>MAB348 | RRID:AB_94882 |

**Table 3.**
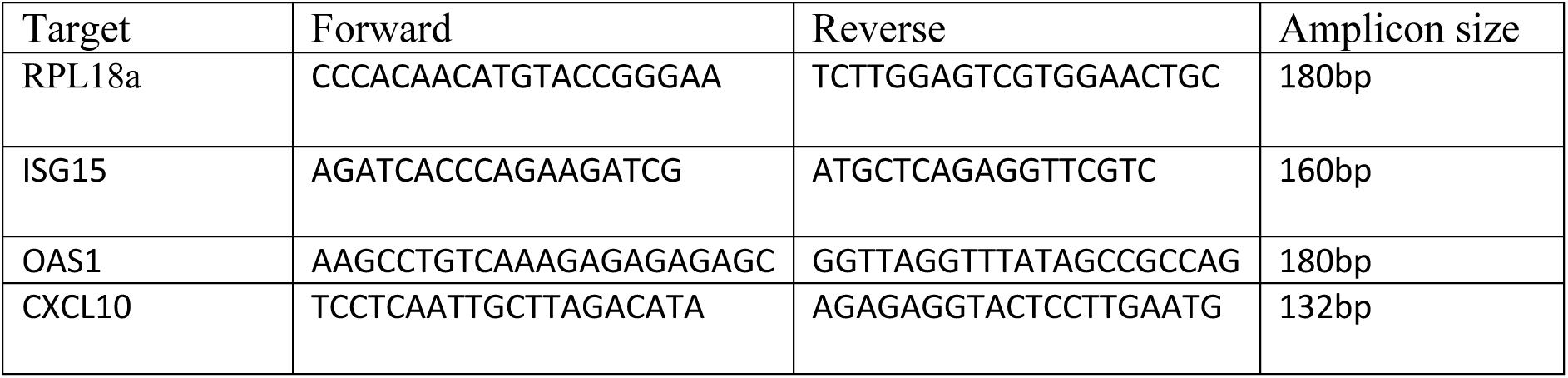
Primers used for qPCR.

We first confirmed the identity of our iPSC-derived astrocyte cultures. Using immunocytochemistry we found homogeneous expression of the astrocyte markers SOX9 and ALDH1L1 (Fig 1A-C). We observed a lower proportion of GFAP immunoreactivity, potentially reflective of lower inflammatory states in these serum-free cultures (Fig 1D) [64]. Transcriptomic analysis confirmed enrichment of astrocyte marker genes in our iPSC-derived astrocyte cultures compared with iPSC-derived neurons (Fig 1E). Principal component analysis revealed separation based on cell type along PC1 and we found that human foetal astrocytes clustered together with our iPSC-astrocytes (Fig 1F). We next examined responses to TIC treatment and confirmed TIC treated samples separated from their untreated counterparts along PC2 (Fig 1F). Cytokine arrays demonstrated the upregulation of inflammatory cytokines and chemokines in astrocyte conditioned media after TIC treatment (Fig S2B).

**Figure 1.**
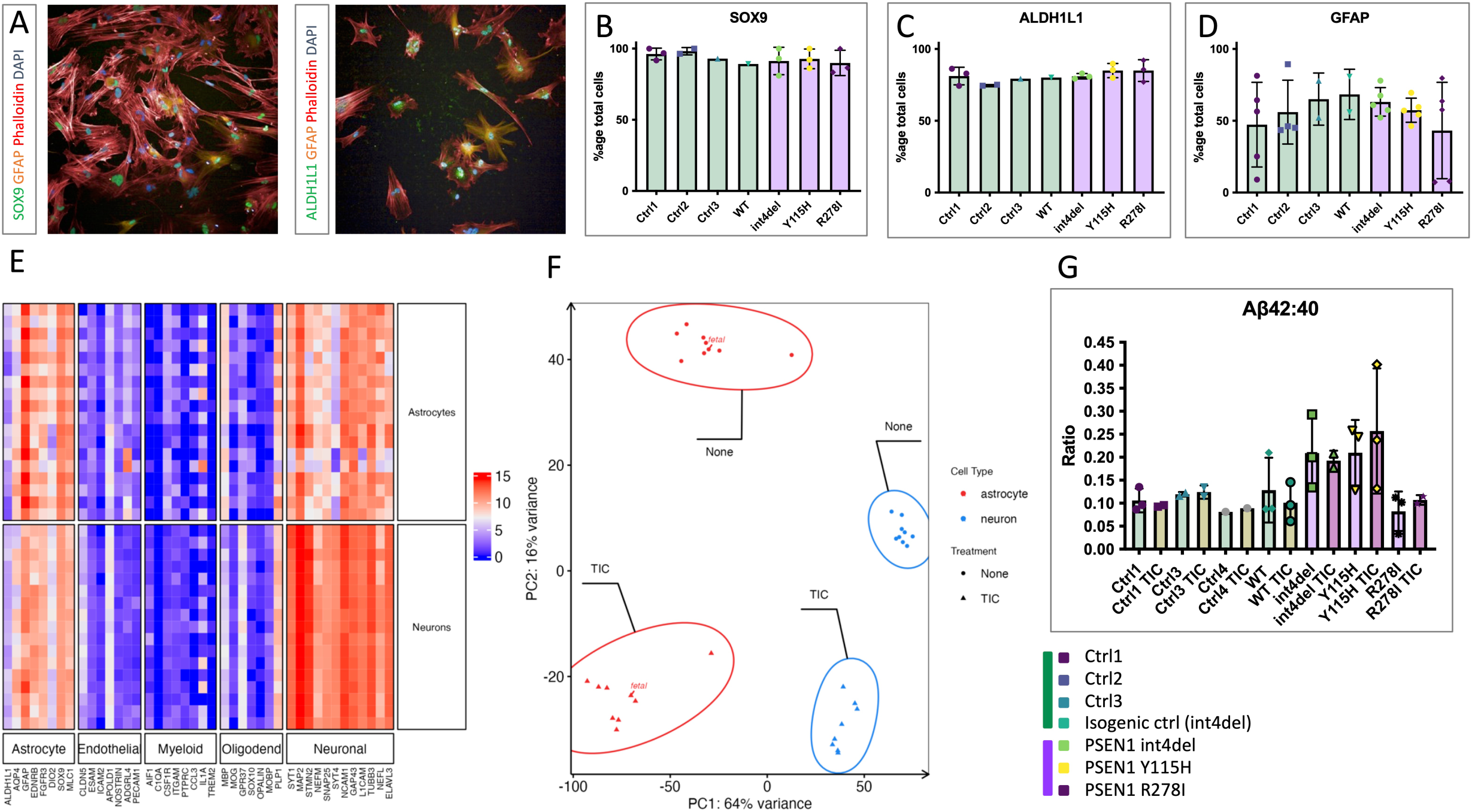
Establishment of an iPSC-derived astrocyte model of familial Alzheimer’s disease. A) Immunocytochemistry of iPSC-astrocytes for markers of astrocytes (SOX9, GFAP, ALDH1L1) and counterstained with phalloidin (f-actin) and DAPI (DNA). B-D) Characterisation of iPSC-astrocyte cultures using high content imaging showing the percentage of cells exhibiting positive immunostaining for astrocyte markers. (data represent 3 independent experimental replicates from 4 control lines and 3 fAD lines total – see Table S1. Technical duplicates are shown for GFAP analysis). E) Transcriptomic characterisation of iPSC-astrocyte cultures in comparison to iPSC-neuronal cultures, showing a relative enrichment of astrocyte markers (data represent 3 independent experimental replicates from 5 control lines and 3 fAD lines total – see Table S1). F) Principal component analysis of transcriptomic data from iPSC neurons, iPSC astrocytes and primary foetal astrocytes, either untreated (none) or stimulated with TNFα, IL1α and C1Q (TIC) for 24 hours. G) ELISA-based quantification of Aβ42 and Aβ40 in iPSC-astrocyte cultures. The ratio of Aβ42:40 is shown, a well-established biomarker of fAD (data represent 3 independent experimental replicates from 4 control lines and 3 fAD lines total – see Table S1).

Importantly, TIC treatment altered neither the Aβ profiles of the astrocyte cultures (Fig 1G) nor measures of cell death (Fig S2C). It should be noted that although Aβ profiles resembled previously described ratios in iPSC-neurons [4], Aβ levels were considerably more variable in astrocyte conditioned media, perhaps reflecting lower amyloidogenic processing (levels of sAPPβ, Fig S2A). Thus, these patient-derived cultures represent a physiological model of fAD astrocytes capable of responding to inflammatory cues.

### PSEN1, PSEN2 and APP are involved in astrocytic responses to inflammatory cues

Based on the literature demonstrating γ-secretase cleavage of cytokine receptors [11, 14], we investigated the role of PSEN1, PSEN2 and APP in the astrocytic response to inflammatory cues. In healthy control astrocytes, *PSEN1* and *PSEN2* gene expression is significantly increased in response to TIC (Fig 2A). This contrasts with iPSC-derived neuronal cultures, which show no significant change in expression (Fig 2B). The upregulation of PSEN1 at the protein level was confirmed by western blotting in iPSC-astrocytes from healthy controls (Fig 2C-D). TIC treatment in *PSEN1* mutant astrocytes led to a non-significant increase in PSEN1. Gene and protein expression of *APP* was not significantly altered by TIC in iPSC-derived astrocytes (Fig 2A); however, we observed a significant accumulation of APP C-terminal fragments upon TIC treatment, representing the substrate for γ-secretase cleavage (Fig 2E-G).

**Figure 2.**
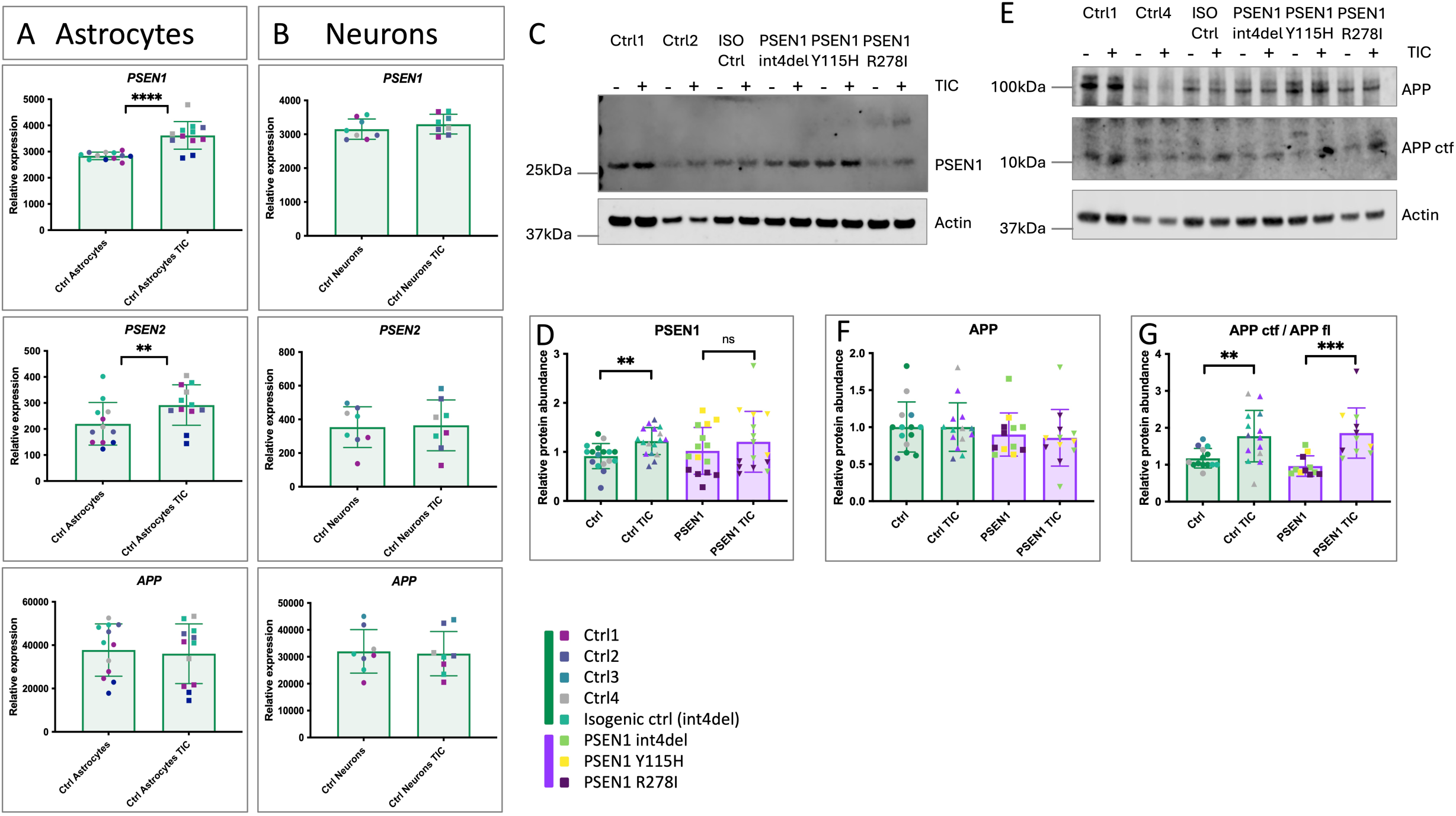
PSEN1, PSEN2 and APP are involved in astrocytic responses to inflammatory cues. *PSEN1* mutations impinge on PSEN1 upregulation. A) Expression of *PSEN1*, *PSEN2* and *APP* in control astrocytes in basal conditions and after 24 hours TNFα, IL1α, C1Q (TIC). Data represent 3 independent experimental replicates from 5 control lines – see Table S1). B) Expression of *PSEN1*, *PSEN2* and *APP* in control neurons in basal conditions and after 24 hours TNFα, IL1α, C1Q (TIC). Data represent 2 independent experimental replicates from 4 control lines – see Table S1). C) Western blotting of iPSC-astrocyte lysates for PSEN1. D) Quantification of PSEN1 Western blotting. Data represent up to 4 batches of astrocytes and up to 5 experimental replicates, including 5 control lines and 3 fAD lines (see Table S1). E) Western blotting of iPSC-astrocyte lysates for APP F-G) Quantification of APP Western blotting. Data represent up to 4 batches of astrocytes and up to 4 experimental replicates, including 5 control lines and 3 *PSEN1* mutant lines (see Table S1). For data separated by each iPSC line, see Fig S7. Pairwise comparisons represent two tailed t-tests, where * = p<0.05, ** = p<0.01, *** = p<0.001, and **** = p<0.0001.

Together, these data suggest that fAD-associated genes are involved in glial responses to inflammation. Moreover, *PSEN1* mutations disrupt the upregulation of PSEN1 at the protein level, potentially impacting its role in inflammatory responses.

### *PSEN1* mutant astrocytes show altered membrane protease gene expression

Regulated intramembrane proteolysis is orchestrated by numerous enzymes. Prior to γ-secretase cleavage, ectodomain shedding of substrates is required to enable loading of the membrane domain into the γ-secretase complex [62]. Shedding is commonly attributed to α-secretases (ADAM10 and ADAM17) or β-secretase (BACE1).

We investigated the expression of γ-secretase, β-secretase and α-secretase components in iPSC-derived astrocytes and the effect of *PSEN1* mutations and inflammatory cues (TIC). As described above, *PSEN1* was upregulated by inflammatory conditions (Fig 2A). Mutations in *PSEN1* did not affect *PSEN1* gene expression itself (Fig 3A). However, *PSEN2* was significantly increased in *PSEN1* mutant astrocytes compared to healthy controls (Fig 3B), possibly reflecting a compensatory response.

**Figure 3.**
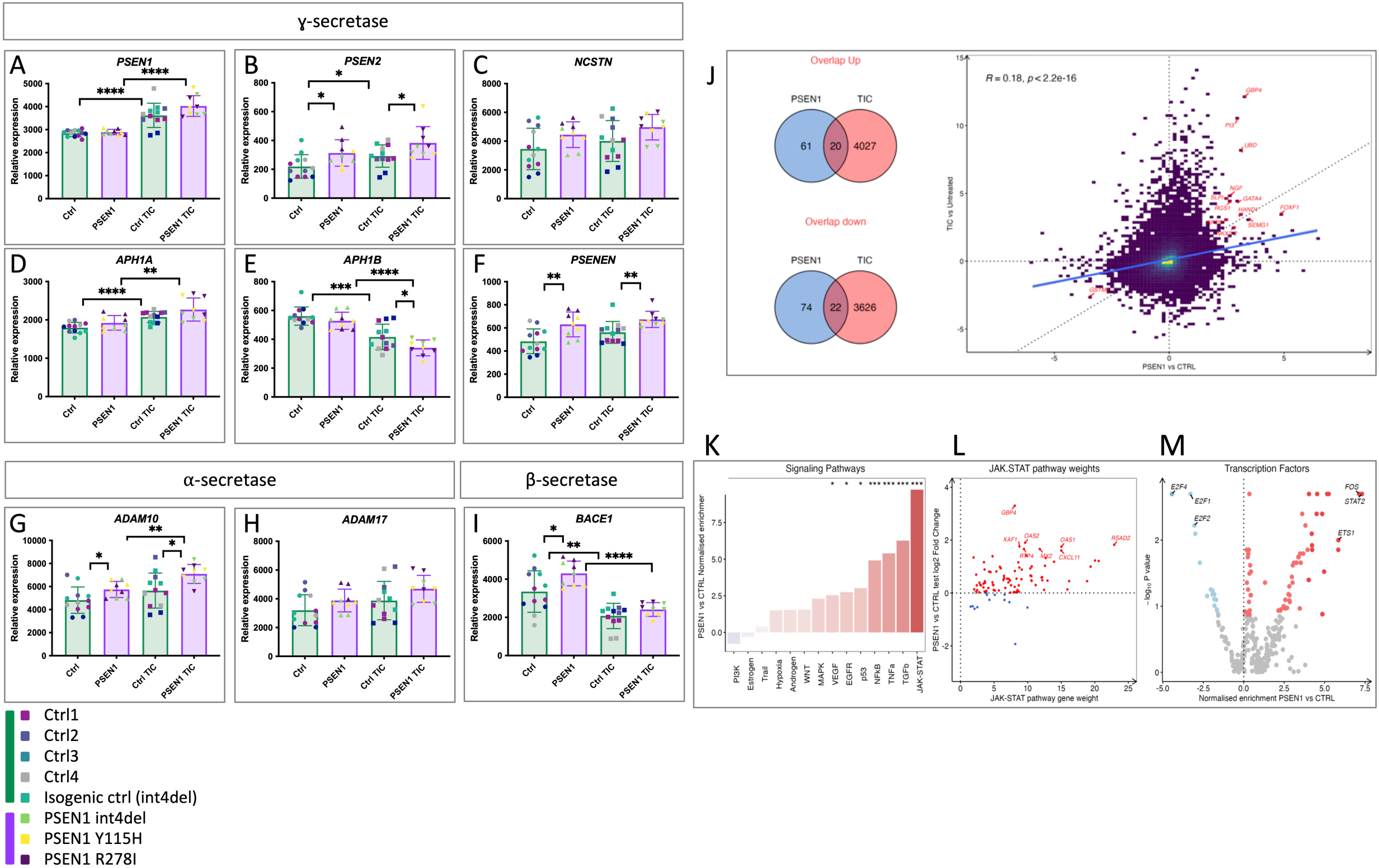
*PSEN1* mutations disrupt regulated intramembrane proteolysis and lead to inflammatory phenotypes. A-I) Expression of γ-secretase, α-secretase and β-secretase components in basal and TIC- treated iPSC-astrocytes, as quantified by transcriptomic analysis. Data represents 3 experimental repeats with 5 controls and 3 *PSEN1* mutant astrocyte lines. J) Correlation analysis of differentially expressed genes (DEGs) in control TIC-treated astrocytes and *PSEN1* mutant untreated astrocytes, relative to control untreated cultures. Of 175 DEGs in *PSEN1* mutant cultures, 42 are shared with TIC treated controls. K-L) Regulon analysis to implicate transcription factors and cell signalling pathways driving differential gene expression in untreated *PSEN1* versus control cultures, analysed via DoRothEA. JAK-STAT2 signalling is highlighted as the most significant pathway. Pairwise comparisons represent two tailed t-tests, where * = p<0.05, ** = p<0.01, *** = p<0.001, and **** = p<0.0001.

In other γ-secretase components, we witnessed upregulated expression in response to TIC with varying degrees of statistical significance (Fig 3C-F) (*APH1B* was downregulated in contrast to its alternate subunit *APH1A*). Interestingly, there was a trend to increased expression of all γ-secretase components at basal levels in *PSEN1* mutant astrocytes compared to healthy controls, mirroring the effect of TIC treatment in healthy control astrocytes. This was most significant in the upregulation of the obligate γ-secretase component *PSENEN*, a subunit for which there is no redundancy (Fig 3F). α-secretase-associated *ADAM10* and *ADAM17* both showed trends towards upregulation in response to inflammation, and *PSEN1* mutations caused significantly upregulated *ADAM10* expression (Fig 3G-H). Expression of the β-secretase gene *BACE1* was reduced by inflammatory stimuli, and we found a significant upregulation in basal conditions in *PSEN1* mutants compared to healthy control patient-derived astrocytes (Fig 3I).

To further investigate altered regulated intramembrane proteolysis, we measured the shed ectodomain of TNFR1, sTNFR1, in astrocyte conditioned media as TNFR1 is a known substrate of γ-secretase [11]. We observed a non-significant trend to increased sTNFR1 in fAD conditioned media in basal conditions (Fig S3A-B). TIC treatment had little effect on TNFR1 shedding.

Together, these data suggest that inflammatory stimuli alter the expression of genes associated with regulated intramembrane proteolysis. In untreated *PSEN1* mutant astrocytes, we observe significantly altered gene expression in a similar direction to the changes seen in response to inflammation, suggesting that inflammatory state may be affected by the presence of mutations in *PSEN1*.

### *PSEN1* mutant astrocytes display augmented inflammatory states

To further investigate the consequence of *PSEN1* mutations on astrocyte function, we performed transcriptomic analysis in untreated and TIC-treated astrocytes.

We found 272 differentially expressed genes (DEGs) in *PSEN1* mutant astrocytes versus healthy control iPSC astrocytes, with 115 upregulated and 157 downregulated DEGs (Fig S4A). Gene Set Enrichment analysis (GSEA, KEGG) showed that DEGs were overrepresented by terms such as cytokine-cytokine receptor (GAGE analysis demonstrates 132 genes with an adjusted P value of 1.1×10^-1^) (Fig S5)[28, 40]. The TNFα pathway was also significantly upregulated (GAGE analysis demonstrates 91 genes with an adjusted P value of 8.5×10^-2^), as was the Alzheimer’s disease term (305 genes with an adjusted P value of 1.4×10^-1^). These data support the hypothesis that *PSEN1* mutations may impinge on regulated intramembrane proteolysis.

We found 2,968 differentially expressed genes in TIC-treated versus untreated healthy control iPSC-astrocytes, with 1746 upregulated and 1222 downregulated in TIC-treated astrocytes (Fig S4B). 3,374 genes were differentially expressed between untreated and TIC-treated astrocytes with the *PSEN1* genotype, with 1,884 upregulated and 1,490 downregulated (Fig S4C). Functional over-representation analysis (KEGG) revealed similar pathways in TIC-treated versus untreated controls for both genotypes. Upregulated pathways included cytokine-cytokine receptor interaction, TNF signalling and NFκB signalling pathways. Downregulated genes did not enrich for KEGG pathways.

When investigating the DEGs compared to control untreated cultures, we observed a significant overlap in *PSEN1* mutant cultures and TIC treated healthy controls (Fig 3J). Of 175 DEGs between untreated controls and untreated *PSEN1* astrocytes, 42 were shared with genes upregulated by TIC in control cultures (up-and downregulated). This suggests similarities between the TIC induced inflammatory phenotype and *PSEN1* mutant astrocytes in basal conditions.

To understand how signalling pathways are activated in *PSEN1* mutant astrocytes under basal conditions, we performed a Signalling Pathway RespOnsive GENes (PROGENy) analysis [7]. This revealed that the most substantial increase in pathway activity in *PSEN1* mutants was in the JAK-STAT pathway, followed by TGFβ, TNFα, and NFκB (Fig 3K). Examining each gene in the JAK-STAT pathway according to its JAK-STAT weighting revealed that the genes with the strongest responsiveness in JAK-STAT activity in *PSEN1* mutant astrocytes included *RSAD2*, *OAS1*, *OAS2*, *CXCL11*, and *GBP4* (Fig 3L). Notably, the JAK-STAT pathway was also substantially upregulated in the TIC response in healthy control astrocytes (Fig S6).

We next inferred the activities of 429 transcription factors (TFs) from their regulon expression within the DoRothEA database. This revealed that STAT2 was the TF with the greatest increase in activity in basal *PSEN1* mutant astrocytes (Fig 3M) followed by FOS and ETS1.

### JAK-STAT2 signalling is activated in *PSEN1* mutant astrocytes

Downstream of ligand (cytokine) binding, phosphorylated tyrosines at the intracellular domain of receptor tyrosine kinases initiate a phosphorylation cascade. Having found JAK-STAT2 as the most upregulated pathway in *PSEN1* mutant astrocytes, we performed western blotting for phosphorylated, activated downstream signalling components including STAT2 and MAPK (p42/p44) (Fig 4A-E).

**Figure 4.**
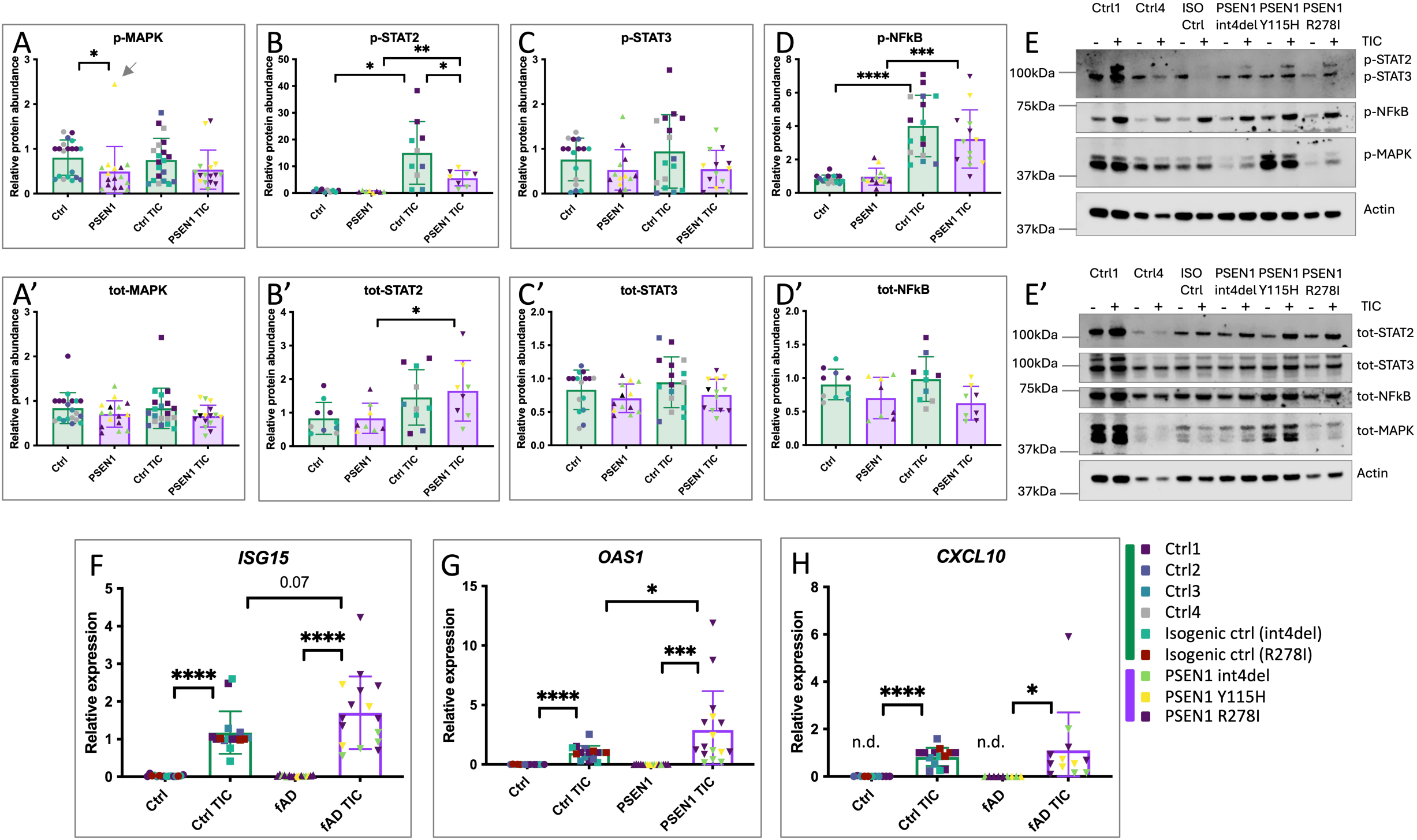
JAK-STAT signalling pathways are disrupted in *PSEN1* mutant astrocytes. A-D) Western blot analysis of total and phosphorylated MAPK (p42/p44), STAT2, STAT3 and NFκB under basal conditions or after 24 hours TIC treatment. Note that one outlier was removed (grey arrow in panel A), due to a z score of 3.47. Data represent up to 4 independent batches and up to 7 technical repeats using 5 controls and 3 *PSEN1* mutant lines (see Table S1). For data separated by iPSC line, see Fig S8. E) Representative Western blot of iPSC-astrocyte lysates with or without TIC treatment for 24 hours. Actin is shown as a loading control. F-H) qPCR analysis of *ISG15*, *OAS1* and *CXCL10*; genes involved in interferon response (associated with JAK-STAT2 signalling). Note that *CXCL10* was rarely detectable in untreated conditions (n.d.). Data represent up to 5 independent batches and up to 6 technical repeats from 6 control lines and 3 *PSEN1* lines, see Table S1. For data separated by iPSC line, see Fig S8. Pairwise comparisons represent two tailed t-tests, where * = p<0.05, ** = p<0.01, *** = p<0.001, and **** = p<0.0001.

We observed a significant reduction in phosphorylated MAPK in *PSEN1* mutant astrocytes under basal conditions (Fig 4A). We also observed a reduced induction of STAT2 phosphorylation after TIC treatment in *PSEN1* cultures compared with healthy controls (Fig 4B). We did not observe significant differences in STAT3 or NFκB signalling (Fig 4C-D). To further validate these data, we quantified JAK-STAT2-associated gene expression by qPCR. We observed significant induction of *ISG15*, *OAS1* and *CXCL10* expression after TIC treatment in control and *PSEN1* mutant astrocytes (Fig 4F-H). We also observed a significantly increased induction of *OAS1* in *PSEN1* astrocytes in response to TIC compared with controls (Fig 4G), supporting data from transcriptomic analyses (e.g., *OAS1* in Fig 3L).

These data support altered phosphorylation of JAK-STAT2 signalling pathway components in response to inflammatory stimuli and further implicate deficits in regulated intramembrane proteolysis of cytokine receptors in *PSEN1* mutant astrocytes.

### Chemical manipulation of γ-secretase reinforces effects of *PSEN1* mutations on JAK-STAT2 signalling

To investigate the effect of γ-secretase activity on inflammatory responses, we treated astrocytes with TIC in combination with a γ-secretase inhibitor (GSI, DAPT) and a γ-secretase modulator (GSM, E2012). γ-secretase modulation has been shown to increase processivity [9, 47], in contrast to γ-secretase inhibition.

Opposing trends were evident in GSM compared with GSI treated cultures. qPCR analyses demonstrated that healthy control astrocytes treated with GSM and TIC had a significantly lower induction of JAK-STAT2 readout genes (*ISG15*, *OAS1* and *CXCL10*) than cultures treated with TIC alone (Fig 5A-C). In contrast, GSI and TIC treated cultures did not significantly differ from TIC treated astrocytes, although the trend was to stronger inflammatory responses with GSI plus TIC compared with TIC alone (Fig 5D-F). Finally, it was interesting to note that GSM treated cultures displayed a higher degree of TNFR1 shedding compared to their untreated counterparts, although this was not the case in cultures with TIC (Fig S3C).

**Figure 5.**
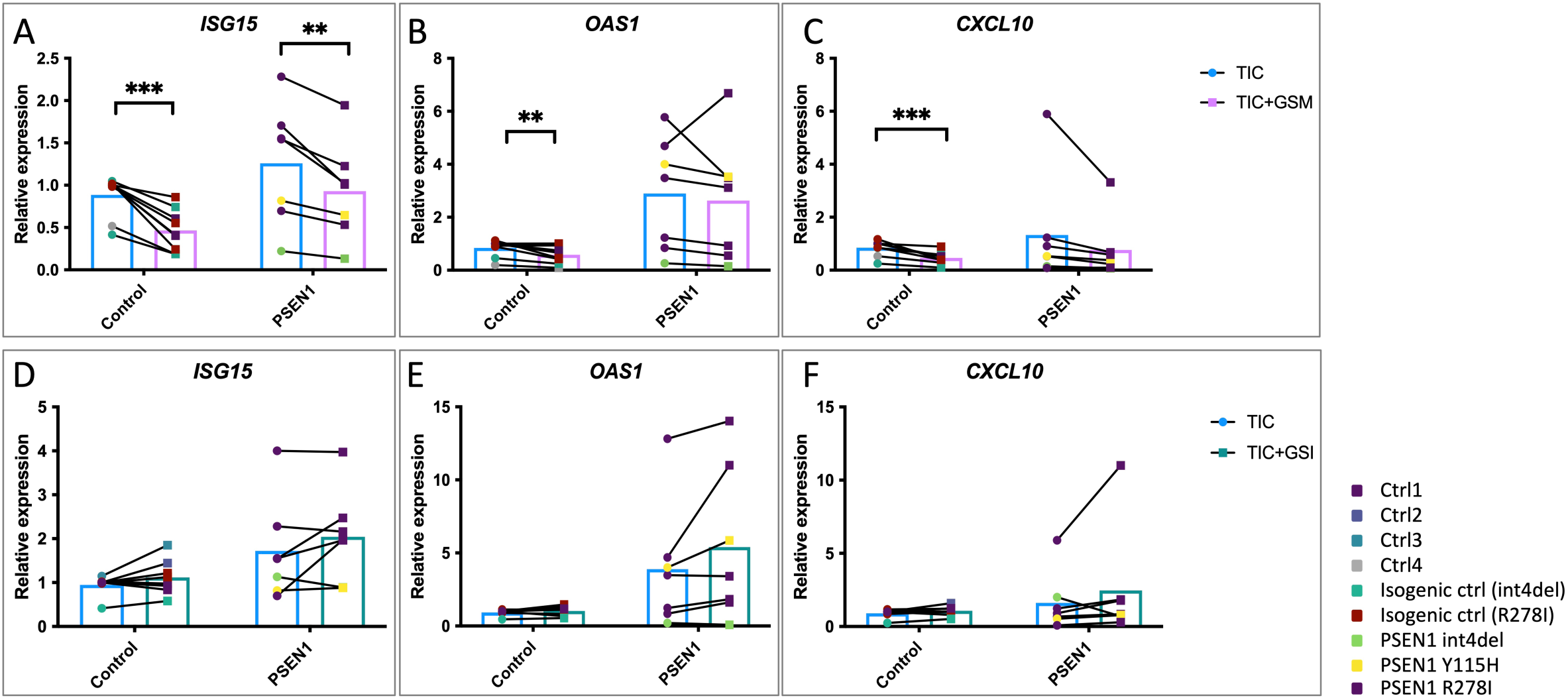
Chemical modulation of γ-secretase reduces interferon-responsive gene expression. A-C) qPCR analysis of interferon responsive gene (*ISG15*, *OAS1* and *CXCL10*) expression after treatment of iPSC-astrocytes with TIC for 24 hours in combination with the γ-secretase modulator (GSM) E2012. Data represent up to 3 independent batches and up to 4 technical repeats from 4 control lines and 3 *PSEN1* lines, see Table S1. D-E) qPCR analysis of interferon responsive gene (*ISG15*, *OAS1* and *CXCL10*) expression after treatment of iPSC-astrocytes with TIC for 24 hours in combination with the γ-secretase inhibitor (GSI) DAPT. Data represent up to 3 independent batches and up to 3 technical repeats from 5 control lines and 3 *PSEN1* lines, see Table S1. Pairwise comparisons represent paired, two tailed t-tests, where * = p<0.05, ** = p<0.01, *** = p<0.001, and **** = p<0.0001.

Together, these data support the finding that manipulating γ-secretase activity has a functional effect on inflammatory responses.

## Discussion

Using patient derived astrocytes, our data support the finding that PSEN1 has a role in the glial response to inflammatory cues. The consequence of fAD-associated mutations in *PSEN1* is altered inflammatory cytokine responses and an enhanced basal inflammatory state in iPSC-derived astrocytes. Together, these data support the notion that *PSEN1* mutations predispose to inflammation.

A predisposition to inflammation would thereby support a two-hit hypothesis in fAD; whereby we propose that *PSEN1* mutations not only impact APP processing but also the cellular response to inflammation, a hypothesis supported by recent work in microglia [26]. Risk factors associated with late onset AD (LOAD) have been shown to link inflammation and disease progression [8, 31], and therefore a two-hit hypothesis of fAD would link disease mechanisms in fAD and LOAD. Indeed, recent work has specifically highlighted JAK-STAT2 and interferon signalling in astrocytes with resilience to AD pathology [32], further reinforcing the findings herein and the potential overlap between pathomechanisms in fAD and LOAD.

Astrocytes have been centrally implicated in Alzheimer’s disease pathobiology previously, such as via the neurotoxic astrocyte phenotype [20, 21, 23, 37]. However, astrocytes are often thought to represent secondary responders to microglial inflammation. The literature supports an early role for astrocyte dysfunction in Alzheimer’s disease [16, 44]. For example, investigations into astrocyte-associated SNPs have linked the cell type with the earliest, amyloid phase of disease, whereas microglial-enriched genes are closely associated with putative downstream tau pathology [67]. The data presented here demonstrate that *PSEN1* mutant astrocytes may respond to disease processes in a cell autonomous manner; i.e., independently from microglia, given that microglia are absent in our cultures and that *PSEN1* mutant astrocytes show predisposition to inflammation in the absence of extrinsic stimuli. In support of this, *in vivo* and *ex vivo* animal studies have shown that reduced astrocyte function is associated with reduced Aβ clearance and increased plaque load [12, 30]. Additionally, recently identified biomarkers in pre-manifest fAD CSF reveal several candidates that are also dysregulated in our fAD patient-derived astrocyte cultures (e.g., SMOC2, STMN2 and SLIT/PDLIM family genes); providing clinical relevance to our model and supporting the presence of astrocyte dysfunction decades before clinical onset [59]. The proinflammatory phenotype of fAD astrocytes is supported by independent iPSC studies which show increased cytokine secretion and reduced neuronal support [45].

Based on our transcriptomic data, we posit that mutations in *PSEN1* broadly impact regulated membrane proteolysis. As a candidate pathway for detailed characterisation, we selected type I interferon signalling; including *OAS1* as a readout which has been identified as an Alzheimer’s disease risk gene [53]. Although interferon signalling has been previously implicated in astrocytes in a disease context [50], it is intriguing that we observe no expression of interferon ligands in these cultures, even after TIC stimulation. Furthermore, we propose that the effects of *PSEN1* mutations may extend to other cell types such as microglia. This is supported by findings demonstrating a role for γ-secretase in microglial state transitions [26] and studies implicating microglial interferon signalling in Down’s syndrome [27].

Our data support an anti-inflammatory role for γ-secretase activity, demonstrated by a decreased interferon response in astrocytes pretreated with γ-secretase modulators (which increase γ-secretase processivity). These findings are relevant to translational efforts, including ongoing γ-secretase modulator trials and the unsuccessful clinical trials of γ-secretase inhibitors, which worsened disease-associated phenotypes concomitant with increased infections rates [13]. Additionally, loss of function mutations in *PSEN1*, *PSENEN* and *NCSTN* all cause familial hidradenitis suppurativa, a chronic inflammatory skin condition [46, 65], further supporting a putative anti-inflammatory role of γ-secretase.

In conclusion, we propose that fAD mutations in *PSEN1* disrupt membrane proteolysis in iPSC-derived astrocytes, impacting the inflammatory state of the cells. Disrupted cytokine signalling may therefore predispose to age-related inflammation in the brain, contributing to the onset and progression of Alzheimer’s disease. This aligns fAD with pathomechanisms of disease in LOAD and could inform rationalised therapeutic intervention.

## Acknowledgements

We would like to acknowledge UCL Genomics for their contribution to the transcriptomics work associated with this investigation. We thank the ICV-iPS (Stéphanie Bigou) and the ICV-Cytometry platforms of ICM, Paris France for editing the PSEN1 R278I clone.

## Supplementary figures

**Figure S1. CRISPR-Cas9 genetic correction of the *PSEN1* R278I mutation in iPSCs.**

A) Sanger sequencing confirms the correction of the point mutation (T>G).

B) Low coverage whole genome sequencing confirms stable karyotype of the isogenic control iPSC line.

**Figure S2. Astrocyte model of familial Alzheimer’s disease.**

A) Western blotting of iPSC-neuron, iPSC-astrocyte, iPSC-microglia and brain lysates for proteins associated with fAD (PSEN1, PSEN2 and APP). GAPDH and Actin serve as loading controls. Lower panel shows conditioned media from the three iPSC cultures, for amyloidogenic processed sAPP (sAPPβ) relative to total shed APP. 3 independent batches are shown from control iPSC lines.

B) Cytokine array using conditioned media (3 batches pooled) from control and *PSEN1* Y115H iPSC-astrocytes, either untreated or treated with TIC for 24 hours. TNFα and IL1α are highlighted in red, as the array may be confounded by recombinant factors added via the TIC treatment.

C) Lactate dehydrogenase assay to analyse cell death in iPSC-astrocyte culture with or without TIC treatment. 3 independent batches are shown for each iPSC line

**Figure S3. Evidence of altered shedding of TNF receptor in PSEN mutant astrocytes and cultures treated with γ-secretase modulator.**

A) ELISA quantification of sTNFR1 in conditioned media normalised to RNA content of the cell pellet.

B) Shows data from A separated by iPSC line.

C) Quantification of sTNFR1 in conditioned media of cultures treated with γ-secretase modulators/inhibitors and in combination with TIC. Normalised to GSI within each group.

**Figure S4. Differential gene expression of astrocytes (genotype and TIC treatment) investigated via transcriptomics.** Volcano plots to represent differentially expressed genes in A) control vs *PSEN1* astrocytes, B) Control untreated versus control TIC-treated astrocytes, C) *PSEN1* untreated versus *PSEN1* TIC-treated astrocytes.

**Figure S5. Evidence of altered cytokine to cytokine receptor interactions via GSEA of *PSEN1* mutant versus healthy control astrocytes under basal conditions**[28, 40].

**Figure S6. DoRothEA analysis of untreated versus TIC treated control astrocytes.** A) Major pathways upregulated by TIC treatment include NFκB, TNFα and JAK-STAT signalling. Transcription factors driving DEGs include *NFKB1*, *RELA* and *STAT2*.

**Figure S7. Data from Fig 2 presented as individual iPSC lines.**

**Figure S8. Data from Fig 4 presented as individual iPSC lines.**

## References

1. Arber C, Belder CRS, Tomczuk F, Gabriele R, Buhidma Y, Farrell C, O’Connor A, Rice H, Lashley T, Fox NC, Ryan NS, Wray S (2024) The presenilin 1 mutation P436S causes familial Alzheimer’s disease with elevated Aβ43 and atypical clinical manifestations. Alzheimer’s & Dementia. doi: 10.1002/alz.13904

2. Arber C, Casey JM, Crawford S, Rambarack N, Yaman U, Wiethoff S, Augustin E, Piers TM, Rostagno A, Ghiso J, Lewis PA, Revesz T, Hardy J, Pocock JM, Houlden H, Schott JM, Salih DA, Lashley T, Wray S (2023) Microglia produce the amyloidogenic ABri peptide in familial British dementia. bioRxiv 1–47. doi: 10.1101/2023.06.27.546552

3. Arber C, Lovejoy C, Harris L, Willumsen N, Alatza A, Casey JM, Lines G, Kerins C, Mueller AK, Zetterberg H, Hardy J, Ryan NS, Fox NC, Lashley T, Wray S (2021) Familial Alzheimer’s Disease Mutations in PSEN1 Lead to Premature Human Stem Cell Neurogenesis. Cell Rep 34:108615. doi: 10.1016/j.celrep.2020.108615

4. Arber C, Toombs J, Lovejoy C, Ryan NS, Paterson RW, Willumsen N, Gkanatsiou E, Portelius E, Blennow K, Heslegrave A, Schott JM, Hardy J, Lashley T, Fox NC, Zetterberg H, Wray S (2020) Familial Alzheimer’s disease patient-derived neurons reveal distinct mutation-specific effects on amyloid beta. Mol Psychiatry 25:2919– 2931. doi: 10.1038/s41380-019-0410-8

5. Arber C, Villegas-Llerena C, Toombs J, Pocock JM, Ryan NS, Fox NC, Zetterberg H, Hardy J, Wray S (2019) Amyloid precursor protein processing in human neurons with an allelic series of the PSEN1 intron 4 deletion mutation and total presenilin-1 knockout. Brain Commun 1:fcz024. doi: 10.1093/braincomms/fcz024

6. Badia-I-Mompel P, Vélez Santiago J, Braunger J, Geiss C, Dimitrov D, Müller-Dott S, Taus P, Dugourd A, Holland CH, Ramirez Flores RO, Saez-Rodriguez J (2022) decoupleR: ensemble of computational methods to infer biological activities from omics data. Bioinformatics advances 2:vbac016. doi: 10.1093/bioadv/vbac016

7. Badia-I-Mompel P, Vélez Santiago J, Braunger J, Geiss C, Dimitrov D, Müller-Dott S, Taus P, Dugourd A, Holland CH, Ramirez Flores RO, Saez-Rodriguez J (2022) decoupleR: ensemble of computational methods to infer biological activities from omics data. Bioinformatics advances 2:vbac016. doi: 10.1093/bioadv/vbac016

8. Bellenguez C, Küçükali F, Jansen IE, Kleineidam L, Moreno-Grau S, Amin N, Naj AC, Campos-Martin R, Grenier-Boley B, Andrade V, Holmans PA, Boland A, Damotte V, van der Lee SJ, Costa MR, Kuulasmaa T, Yang Q, de Rojas I, Bis JC, Yaqub A, Prokic I, Chapuis J, Ahmad S, Giedraitis V, Aarsland D, Garcia-Gonzalez P, Abdelnour C, Alarcón-Martín E, Alcolea D, Alegret M, Alvarez I, Álvarez V, Armstrong NJ, Tsolaki A, Antúnez C, Appollonio I, Arcaro M, Archetti S, Pastor AA, Arosio B, Athanasiu L, Bailly H, Banaj N, Baquero M, Barral S, Beiser A, Pastor AB, Below JE, Benchek P, Benussi L, Berr C, Besse C, Bessi V, Binetti G, Bizarro A, Blesa R, Boada M, Boerwinkle E, Borroni B, Boschi S, Bossù P, Bråthen G, Bressler J, Bresner C, Brodaty H, Brookes KJ, Brusco LI, Buiza-Rueda D, Bûrger K, Burholt V, Bush WS, Calero M, Cantwell LB, Chene G, Chung J, Cuccaro ML, Carracedo Á, Cecchetti R, Cervera-Carles L, Charbonnier C, Chen H-H, Chillotti C, Ciccone S, Claassen JAHR, Clark C, Conti E, Corma-Gómez A, Costantini E, Custodero C, Daian D, Dalmasso MC, Daniele A, Dardiotis E, Dartigues J-F, de Deyn PP, de Paiva Lopes K, de Witte LD, Debette S, Deckert J, del Ser T, Denning N, DeStefano A, Dichgans M, Diehl-Schmid J, Diez-Fairen M, Rossi PD, Djurovic S, Duron E, Düzel E, Dufouil C, Eiriksdottir G, Engelborghs S, Escott-Price V, Espinosa A, Ewers M, Faber KM, Fabrizio T, Nielsen SF, Fardo DW, Farotti L, Fenoglio C, Fernández-Fuertes M, Ferrari R, Ferreira CB, Ferri E, Fin B, Fischer P, Fladby T, Fließbach K, Fongang B, Fornage M, Fortea J, Foroud TM, Fostinelli S, Fox NC, Franco-Macías E, Bullido MJ, Frank-García A, Froelich L, Fulton-Howard B, Galimberti D, García-Alberca JM, García-González P, Garcia-Madrona S, Garcia-Ribas G, Ghidoni R, Giegling I, Giorgio G, Goate AM, Goldhardt O, Gomez-Fonseca D, González-Pérez A, Graff C, Grande G, Green E, Grimmer T, Grünblatt E, Grunin M, Gudnason V, Guetta-Baranes T, Haapasalo A, Hadjigeorgiou G, Haines JL, Hamilton-Nelson KL, Hampel H, Hanon O, Hardy J, Hartmann AM, Hausner L, Harwood J, Heilmann-Heimbach S, Helisalmi S, Heneka MT, Hernández I, Herrmann MJ, Hoffmann P, Holmes C, Holstege H, Vilas RH, Hulsman M, Humphrey J, Biessels GJ, Jian X, Johansson C, Jun GR, Kastumata Y, Kauwe J, Kehoe PG, Kilander L, Ståhlbom AK, Kivipelto M, Koivisto A, Kornhuber J, Kosmidis MH, Kukull WA, Kuksa PP, Kunkle BW, Kuzma AB, Lage C, Laukka EJ, Launer L, Lauria A, Lee C-Y, Lehtisalo J, Lerch O, Lleó A, Longstreth W, Lopez O, de Munain AL, Love S, Löwemark M, Luckcuck L, Lunetta KL, Ma Y, Macías J, MacLeod CA, Maier W, Mangialasche F, Spallazzi M, Marquié M, Marshall R, Martin ER, Montes AM, Rodríguez CM, Masullo C, Mayeux R, Mead S, Mecocci P, Medina M, Meggy A, Mehrabian S, Mendoza S, Menéndez-González M, Mir P, Moebus S, Mol M, Molina-Porcel L, Montrreal L, Morelli L, Moreno F, Morgan K, Mosley T, Nöthen MM, Muchnik C, Mukherjee S, Nacmias B, Ngandu T, Nicolas G, Nordestgaard BG, Olaso R, Orellana A, Orsini M, Ortega G, Padovani A, Paolo C, Papenberg G, Parnetti L, Pasquier F, Pastor P, Peloso G, Pérez-Cordón A, Pérez-Tur J, Pericard P, Peters O, Pijnenburg YAL, Pineda JA, Piñol-Ripoll G, Pisanu C, Polak T, Popp J, Posthuma D, Priller J, Puerta R, Quenez O, Quintela I, Thomassen JQ, Rábano A, Rainero I, Rajabli F, Ramakers I, Real LM, Reinders MJT, Reitz C, Reyes-Dumeyer D, Ridge P, Riedel-Heller S, Riederer P, Roberto N, Rodriguez-Rodriguez E, Rongve A, Allende IR, Rosende-Roca M, Royo JL, Rubino E, Rujescu D, Sáez ME, Sakka P, Saltvedt I, Sanabria Á, Sánchez-Arjona MB, Sanchez-Garcia F, Juan PS, Sánchez-Valle R, Sando SB, Sarnowski C, Satizabal CL, Scamosci M, Scarmeas N, Scarpini E, Scheltens P, Scherbaum N, Scherer M, Schmid M, Schneider A, Schott JM, Selbæk G, Seripa D, Serrano M, Sha J, Shadrin AA, Skrobot O, Slifer S, Snijders GJL, Soininen H, Solfrizzi V, Solomon A, Song Y, Sorbi S, Sotolongo-Grau O, Spalletta G, Spottke A, Squassina A, Stordal E, Tartan JP, Tárraga L, Tesí N, Thalamuthu A, Thomas T, Tosto G, Traykov L, Tremolizzo L, Tybjærg-Hansen A, Uitterlinden A, Ullgren A, Ulstein I, Valero S, Valladares O, Broeckhoven C Van, Vance J, Vardarajan BN, van der Lugt A, Dongen J Van, van Rooij J, van Swieten J, Vandenberghe R, Verhey F, Vidal J-S, Vogelgsang J, Vyhnalek M, Wagner M, Wallon D, Wang L-S, Wang R, Weinhold L, Wiltfang J, Windle G, Woods B, Yannakoulia M, Zare H, Zhao Y, Zhang X, Zhu C, Zulaica M, Laczo J, Matoska V, Serpente M, Assogna F, Piras F, Piras F, Ciullo V, Shofany J, Ferrarese C, Andreoni S, Sala G, Zoia CP, Zompo M Del, Benussi A, Bastiani P, Takalo M, Natunen T, Laatikainen T, Tuomilehto J, Antikainen R, Strandberg T, Lindström J, Peltonen M, Abraham R, Al-Chalabi A, Bass NJ, Brayne C, Brown KS, Collinge J, Craig D, Deloukas P, Fox N, Gerrish A, Gill M, Gwilliam R, Harold D, Hollingworth P, Johnston JA, Jones L, Lawlor B, Livingston G, Lovestone S, Lupton M, Lynch A, Mann D, McGuinness B, McQuillin A, O’Donovan MC, Owen MJ, Passmore P, Powell JF, Proitsi P, Rossor M, Shaw CE, Smith AD, Gurling H, Todd S, Mummery C, Ryan N, Lacidogna G, Adarmes-Gómez A, Mauleón A, Pancho A, Gailhajenet A, Lafuente A, Macias-García D, Martín E, Pelejà E, Carrillo F, Merlín IS, Garrote-Espina L, Vargas L, Carrion-Claro M, Marín M, Labrador M, Buendia M, Alonso MD, Guitart M, Moreno M, Ibarria M, Periñán M, Aguilera N, Gómez-Garre P, Cañabate P, Escuela R, Pineda-Sánchez R, Vigo-Ortega R, Jesús S, Preckler S, Rodrigo-Herrero S, Diego S, Vacca A, Roveta F, Salvadori N, Chipi E, Boecker H, Laske C, Perneczky R, Anastasiou C, Janowitz D, Malik R, Anastasiou A, Parveen K, Lage C, López-García S, Antonell A, Mihova KY, Belezhanska D, Weber H, Kochen S, Solis P, Medel N, Lisso J, Sevillano Z, Politis DG, Cores V, Cuesta C, Ortiz C, Bacha JI, Rios M, Saenz A, Abalos MS, Kohler E, Palacio DL, Etchepareborda I, Kohler M, Novack G, Prestia FA, Galeano P, Castaño EM, Germani S, Toso CR, Rojo M, Ingino C, Mangone C, Rubinsztein DC, Teipel S, Fievet N, Deramerourt V, Forsell C, Thonberg H, Bjerke M, Roeck E De, Martínez-Larrad MT, Olivar N, Aguilera N, Cano A, Cañabate P, Macias J, Maroñas O, Nuñez-Llaves R, Olivé C, Pelejá E, Adarmes-Gómez AD, Alonso MD, Amer-Ferrer G, Antequera M, Burguera JA, Carrillo F, Carrión-Claro M, Casajeros MJ, Martinez de Pancorbo M, Escuela R, Garrote-Espina L, Gómez-Garre P, Hevilla S, Jesús S, Espinosa MAL, Legaz A, López-García S, Macias-García D, Manzanares S, Marín M, Marín-Muñoz J, Marín T, Martínez B, Martínez V, Martínez-Lage Álvarez P, Iriarte MM, Periñán-Tocino MT, Pineda-Sánchez R, Real de Asúa D, Rodrigo S, Sastre I, Vicente MP, Vigo-Ortega R, Vivancos L, Epelbaum J, Hannequin D, campion D, Deramecourt V, Tzourio C, Brice A, Dubois B, Williams A, Thomas C, Davies C, Nash W, Dowzell K, Morales AC, Bernardo-Harrington M, Turton J, Lord J, Brown K, Vardy E, Fisher E, Warren JD, Rossor M, Ryan NS, Guerreiro R, Uphill J, Bass N, Heun R, Kölsch H, Schürmann B, Lacour A, Herold C, Johnston JA, Passmore P, Powell J, Patel Y, Hodges A, Becker T, Warden D, Wilcock G, Clarke R, Deloukas P, Ben-Shlomo Y, Hooper NM, Pickering-Brown S, Sussams R, Warner N, Bayer A, Heuser I, Drichel D, Klopp N, Mayhaus M, Riemenschneider M, Pinchler S, Feulner T, Gu W, van den Bussche H, Hüll M, Frölich L, Wichmann H-E, Jöckel K-H, O’Donovan M, Owen M, Bahrami S, Bosnes I, Selnes P, Bergh S, Palotie A, Daly M, Jacob H, Matakidou A, Runz H, John S, Plenge R, McCarthy M, Hunkapiller J, Ehm M, Waterworth D, Fox C, Malarstig A, Klinger K, Call K, Behrens T, Loerch P, Mäkelä T, Kaprio J, Virolainen P, Pulkki K, Kilpi T, Perola M, Partanen J, Pitkäranta A, Kaarteenaho R, Vainio S, Turpeinen M, Serpi R, Laitinen T, Mäkelä J, Kosma V-M, Kujala U, Tuovila O, Hendolin M, Pakkanen R, Waring J, Riley-Gillis B, Liu J, Biswas S, Diogo D, Marshall C, Hu X, Gossel M, Graham R, Cummings B, Ripatti S, Schleutker J, Arvas M, Carpén O, Hinttala R, Kettunen J, Mannermaa A, Laukkanen J, Julkunen V, Remes A, Kälviäinen R, Peltola J, Tienari P, Rinne J, Ziemann A, Waring J, Esmaeeli S, Smaoui N, Lehtonen A, Eaton S, Lahdenperä S, van Adelsberg J, Michon J, Kerchner G, Bowers N, Teng E, Eicher J, Mehta V, Gormley P, Linden K, Whelan C, Xu F, Pulford D, Färkkilä M, Pikkarainen S, Jussila A, Blomster T, Kiviniemi M, Voutilainen M, Georgantas B, Heap G, Rahimov F, Usiskin K, Lu T, Oh D, Kalpala K, Miller M, McCarthy L, Eklund K, Palomäki A, Isomäki P, Pirilä L, Kaipiainen-Seppänen O, Huhtakangas J, Lertratanakul A, Hochfeld M, Bing N, Gordillo JE, Mars N, Pelkonen M, Kauppi P, Kankaanranta H, Harju T, Close D, Greenberg S, Chen H, Betts J, Ghosh S, Salomaa V, Niiranen T, Juonala M, Metsärinne K, Kähönen M, Junttila J, Laakso M, Pihlajamäki J, Sinisalo J, Taskinen M-R, Tuomi T, Challis B, Peterson A, Chu A, Parkkinen J, Muslin A, Joensuu H, Meretoja T, Aaltonen L, Mattson J, Auranen A, Karihtala P, Kauppila S, Auvinen P, Elenius K, Popovic R, Schutzman J, Loboda A, Chhibber A, Lehtonen H, McDonough S, Crohns M, Kulkarni D, Kaarniranta K, Turunen JA, Ollila T, Seitsonen S, Uusitalo H, Aaltonen V, Uusitalo-Järvinen H, Luodonpää M, Hautala N, Loomis S, Strauss E, Chen H, Podgornaia A, Hoffman J, Tasanen K, Huilaja L, Hannula-Jouppi K, Salmi T, Peltonen S, Koulu L, Harvima I, Wu Y, Choy D, Pussinen P, Salminen A, Salo T, Rice D, Nieminen P, Palotie U, Siponen M, Suominen L, Mäntylä P, Gursoy U, Anttonen V, Sipilä K, Davis JW, Quarless D, Petrovski S, Wigmore E, Chen C-Y, Bronson P, Tsai E, Huang Y, Maranville J, Shaikho E, Mohammed E, Wadhawan S, Kvikstad E, Caliskan M, Chang D, Bhangale T, Pendergrass S, Holzinger E, Chen X, Hedman Å, King KS, Wang C, Xu E, Auge F, Chatelain C, Rajpal D, Liu D, Call K, Xia T, Brauer M, Kurki M, Karjalainen J, Havulinna A, Jalanko A, Palta P, della Briotta Parolo P, Zhou W, Lemmelä S, Rivas M, Harju J, Lehisto A, Ganna A, Llorens V, Laivuori H, Rüeger S, Niemi ME, Tukiainen T, Reeve MP, Heyne H, Palin K, Garcia-Tabuenca J, Siirtola H, Kiiskinen T, Lee J, Tsuo K, Elliott A, Kristiansson K, Hyvärinen K, Ritari J, Koskinen M, Pylkäs K, Kalaoja M, Karjalainen M, Mantere T, Kangasniemi E, Heikkinen S, Laakkonen E, Sipeky C, Heron S, Karlsson A, Jambulingam D, Rathinakannan VS, Kajanne R, Aavikko M, Jiménez MG, della Briotta Parola P, Lehistö A, Kanai M, Kaunisto M, Kilpeläinen E, Sipilä TP, Brein G, Awaisa G, Shcherban A, Donner K, Loukola A, Laiho P, Sistonen T, Kaiharju E, Laukkanen M, Järvensivu E, Lähteenmäki S, Männikkö L, Wong R, Mattsson H, Hiekkalinna T, Paajanen T, Pärn K, Gracia-Tabuenca J, Abner E, Adams PM, Aguirre A, Albert MS, Albin RL, Allen M, Alvarez L, Apostolova LG, Arnold SE, Asthana S, Atwood CS, Ayres G, Baldwin CT, Barber RC, Barnes LL, Barral S, Beach TG, Becker JT, Beecham GW, Beekly D, Below JE, Benchek P, Benitez BA, Bennett D, Bertelson J, Margaret FE, Bird TD, Blacker D, Boeve BF, Bowen JD, Boxer A, Brewer J, Burke JR, Burns JM, Bush WS, Buxbaum JD, Cairns NJ, Cao C, Carlson CS, Carlsson CM, Carney RM, Carrasquillo MM, Chasse S, Chesselet M-F, Chesi A, Chin NA, Chui HC, Chung J, Craft S, Crane PK, Cribbs DH, Crocco EA, Cruchaga C, Cuccaro ML, Cullum M, Darby E, Davis B, De Jager PL, DeCarli C, DeToledo J, Dick M, Dickson DW, Dombroski BA, Doody RS, Duara R, Ertekin-Taner N, Evans DA, Fairchild TJ, Fallon KB, Farlow MR, Farrell JJ, Fernandez-Hernandez V, Ferris S, Frosch MP, Fulton-Howard B, Galasko DR, Gamboa A, Gearing M, Geschwind DH, Ghetti B, Gilbert JR, Grabowski TJ, Graff-Radford NR, Grant SFA, Green RC, Growdon JH, Haines JL, Hakonarson H, Hall J, Hamilton RL, Harari O, Harrell LE, Haut J, Head E, Henderson VW, Hernandez M, Hohman T, Honig LS, Huebinger RM, Huentelman MJ, Hulette CM, Hyman BT, Hynan LS, Ibanez L, Jarvik GP, Jayadev S, Jin L-W, Johnson K, Johnson L, Kamboh MI, Karydas AM, Katz MJ, Kaye JA, Keene CD, Khaleeq A, Kim R, Knebl J, Kowall NW, Kramer JH, Kuksa PP, LaFerla FM, Lah JJ, Larson EB, Lee C-Y, Lee EB, Lerner A, Leung YY, Leverenz JB, Levey AI, Li M, Lieberman AP, Lipton RB, Logue M, Lyketsos CG, Malamon J, Mains D, Marson DC, Martiniuk F, Mash DC, Masliah E, Massman P, Masurkar A, McCormick WC, McCurry SM, McDavid AN, McDonough S, McKee AC, Mesulam M, Mez J, Miller BL, Miller CA, Miller JW, Montine TJ, Monuki ES, Morris JC, Myers AJ, Nguyen T, O’Bryant S, Olichney JM, Ory M, Palmer R, Parisi JE, Paulson HL, Pavlik V, Paydarfar D, Perez V, Peskind E, Petersen RC, Phillips-Cremins JE, Pierce A, Polk M, Poon WW, Potter H, Qu L, Quiceno M, Quinn JF, Raj A, Raskind M, Reiman EM, Reisberg B, Reisch JS, Ringman JM, Roberson ED, Rodriguear M, Rogaeva E, Rosen HJ, Rosenberg RN, Royall DR, Sager MA, Sano M, Saykin AJ, Schneider JA, Schneider LS, Seeley WW, Slifer SH, Small S, Smith AG, Smith JP, Song YE, Sonnen JA, Spina S, George-Hyslop PS, Stern RA, Stevens AB, Strittmatter SM, Sultzer D, Swerdlow RH, Tanzi RE, Tilson JL, Trojanowski JQ, Troncoso JC, Tsuang DW, Valladares O, Van Deerlin VM, van Eldik LJ, Vassar R, Vinters H V., Vonsattel J-P, Weintraub S, Welsh-Bohmer KA, Whitehead PL, Wijsman EM, Wilhelmsen KC, Williams B, Williamson J, Wilms H, Wingo TS, Wisniewski T, Woltjer RL, Woon M, Wright CB, Wu C-K, Younkin SG, Yu C-E, Yu L, Zhang Y, Zhao Y, Zhu X, Adams H, Akinyemi RO, Ali M, Armstrong N, Aparicio HJ, Bahadori M, Becker JT, Breteler M, Chasman D, Chauhan G, Comic H, Cox S, Cupples AL, Davies G, DeCarli CS, Duperron M-G, Dupuis J, Evans T, Fan F, Fitzpatrick A, Fohner AE, Ganguli M, Geerlings M, Glatt SJ, Gonzalez HM, Goss M, Grabe H, Habes M, Heckbert SR, Hofer E, Hong E, Hughes T, Kautz TF, Knol M, Kremen W, Lacaze P, Lahti J, Grand Q Le, Litkowski E, Li S, Liu D, Liu X, Loitfelder M, Manning A, Maillard P, Marioni R, Mazoyer B, van Lent DM, Mei H, Mishra A, Nyquist P, O’Connell J, Patel Y, Paus T, Pausova Z, Raikkonen-Talvitie K, Riaz M, Rich S, Rotter J, Romero J, Roshchupkin G, Saba Y, Sargurupremraj M, Schmidt H, Schmidt R, Shulman JM, Smith J, Sekhar H, Rajula R, Shin J, Simino J, Sliz E, Teumer A, Thomas A, Tin A, Tucker-Drob E, Vojinovic D, Wang Y, Weinstein G, Williams D, Wittfeld K, Yanek L, Yang Y, Farrer LA, Psaty BM, Ghanbari M, Raj T, Sachdev P, Mather K, Jessen F, Ikram MA, de Mendonça A, Hort J, Tsolaki M, Pericak-Vance MA, Amouyel P, Williams J, Frikke-Schmidt R, Clarimon J, Deleuze J-F, Rossi G, Seshadri S, Andreassen OA, Ingelsson M, Hiltunen M, Sleegers K, Schellenberg GD, van Duijn CM, Sims R, van der Flier WM, Ruiz A, Ramirez A, Lambert J-C (2022) New insights into the genetic etiology of Alzheimer’s disease and related dementias. Nat Genet 54:412–436. doi: 10.1038/s41588-022-01024-z

9. Borgegard T, Juréus A, Olsson F, Rosqvist S, Sabirsh A, Rotticci D, Paulsen K, Klintenberg R, Yan H, Waldman M, Stromberg K, Nord J, Johansson J, Regner A, Parpal S, Malinowsky D, Radesater AC, Li T, Singh R, Eriksson H, Lundkvist J (2012) First and second generation γ-secretase modulators (GSMs) modulate amyloid-β (Aβ) peptide production through different mechanisms. Journal of Biological Chemistry 287:11810–11819. doi: 10.1074/jbc.M111.305227

10. Bruntraeger M, Byrne M, Long K, Bassett AR (2019) Editing the Genome of Human Induced Pluripotent Stem Cells Using CRISPR/Cas9 Ribonucleoprotein Complexes. Methods Mol Biol 1961:153–183. doi: 10.1007/978-1-4939-9170-9_11

11. Chhibber-Goel J, Coleman-Vaughan C, Agrawal V, Sawhney N, Hickey E, Powell JC, McCarthy J V. (2016) γ-Secretase Activity Is Required for Regulated Intramembrane Proteolysis of Tumor Necrosis Factor (TNF) Receptor 1 and TNF-mediated Pro-apoptotic Signaling. Journal of Biological Chemistry 291:5971–5985. doi: 10.1074/jbc.M115.679076

12. Davis N, Mota BC, Stead L, Palmer EO, Lombardero L, Rodriguez-Puertas R, de Paola V, Barnes SJ, Sastre M (2020) Ablation of astrocytes affects Aβ degradation, microglia activation and synaptic connectivity in an ex vivo model of Alzheimer’s disease. Alzheimer’s & Dementia 16:1–12. doi: 10.1002/alz.047419

13. Doody RS, Raman R, Farlow M, Iwatsubo T, Vellas B, Joffe S, Kieburtz K, He F, Sun X, Thomas RG, Aisen PS, Siemers E, Sethuraman G, Mohs R (2013) A Phase 3 Trial of Semagacestat for Treatment of Alzheimer’s Disease. New England Journal of Medicine 369:341–350. doi: 10.1056/NEJMoa1210951

14. Elzinga BM, Twomey C, Powell JC, Harte F, McCarthy J V. (2009) Interleukin-1 Receptor Type 1 Is a Substrate for γ-Secretase-dependent Regulated Intramembrane Proteolysis. Journal of Biological Chemistry 284:1394–1409. doi: 10.1074/jbc.M803108200

15. Ewels PA, Peltzer A, Fillinger S, Patel H, Alneberg J, Wilm A, Garcia MU, Di Tommaso P, Nahnsen S (2020) The nf-core framework for community-curated bioinformatics pipelines. Nat Biotechnol 38:276–278. doi: 10.1038/s41587-020-0439-x

16. Fontana IC, Scarpa M, Malarte ML, Rocha FM, Ausellé-Bosch S, Bluma M, Bucci M, Chiotis K, Kumar A, Nordberg A (2023) Astrocyte Signature in Alzheimer’s Disease Continuum through a Multi-PET Tracer Imaging Perspective. Cells 12:1–12. doi: 10.3390/cells12111469

17. Garcia-Reitboeck P, Phillips A, Piers TM, Villegas-Llerena C, Butler M, Mallach A, Rodrigues C, Arber CE, Heslegrave A, Zetterberg H, Neumann H, Neame S, Houlden H, Hardy J, Pocock JM (2018) Human Induced Pluripotent Stem Cell-Derived Microglia-Like Cells Harboring TREM2 Missense Mutations Show Specific Deficits in Phagocytosis. Cell Rep 24:2300–2311. doi: 10.1016/j.celrep.2018.07.094

18. Goate A, Chartier-Harlin M-C, Mullan M, Brown J, Crawford F, Fidani L, Giuffra L, Haynes A, Irving N, James L, Mant R, Newton P, Rooke K, Roques P, Talbot C, Pericak-Vance M, Roses A, Williamson R, Rossor M, Owen M, Hardy J (1991) Segregation of a missense mutation in the amyloid precursor protein gene with familial Alzheimer’s disease. Nature 349:704–706. doi: 10.1038/349704a0

19. Güner G, Lichtenthaler SF (2020) The substrate repertoire of γ-secretase/presenilin. Semin Cell Dev Biol 105:27–42. doi: 10.1016/j.semcdb.2020.05.019

20. Guttenplan KA, Weigel MK, Adler DI, Couthouis J, Liddelow SA, Gitler AD, Barres BA (2020) Knockout of reactive astrocyte activating factors slows disease progression in an ALS mouse model. Nat Commun 11:3753. doi: 10.1038/s41467-020-17514-9

21. Guttenplan KA, Weigel MK, Prakash P, Wijewardhane PR, Hasel P, Rufen-Blanchette U, Münch AE, Blum JA, Fine J, Neal MC, Bruce KD, Gitler AD, Chopra G, Liddelow SA, Barres BA (2021) Neurotoxic reactive astrocytes induce cell death via saturated lipids. Nature 599:102–107. doi: 10.1038/s41586-021-03960-y

22. Haapasalo A, Kovacs DM (2011) The many substrates of presenilin/γ-secretase. Journal of Alzheimer’s Disease 25:3–28. doi: 10.3233/JAD-2011-101065

23. Habib N, McCabe C, Medina S, Varshavsky M, Kitsberg D, Dvir-Szternfeld R, Green G, Dionne D, Nguyen L, Marshall JL, Chen F, Zhang F, Kaplan T, Regev A, Schwartz M (2020) Disease-associated astrocytes in Alzheimer’s disease and aging. Nat Neurosci 23:701–706. doi: 10.1038/s41593-020-0624-8

24. Hall CE, Yao Z, Choi M, Tyzack GE, Serio A, Luisier R, Harley J, Preza E, Arber C, Crisp SJ, Watson PMD, Kullmann DM, Abramov AY, Wray S, Burley R, Loh SHY, Martins LM, Stevens MM, Luscombe NM, Sibley CR, Lakatos A, Ule J, Gandhi S, Patani R (2017) Progressive Motor Neuron Pathology and the Role of Astrocytes in a Human Stem Cell Model of VCP-Related ALS. Cell Rep 19:1739–1749. doi: 10.1016/j.celrep.2017.05.024

25. Hardy JA, Higgins GA (1992) Alzheimer’s Disease: The Amyloid Cascade Hypothesis. Science (1979) 256:184–185. doi: 10.1126/science.1566067

26. Hou P, Zielonka M, Serneels L, Martinez-Muriana A, Fattorelli N, Wolfs L, Poovathingal S, T’Syen D, Balusu S, Theys T, Fiers M, Mancuso R, Howden AJM, De Strooper B (2023) The γ-secretase substrate proteome and its role in cell signaling regulation. Mol Cell 83:4106–4122.e10. doi: 10.1016/j.molcel.2023.10.029

27. Jin M, Xu R, Wang L, Alam MM, Ma Z, Zhu S, Martini AC, Jadali A, Bernabucci M, Xie P, Kwan KY, Pang ZP, Head E, Liu Y, Hart RP, Jiang P (2022) Type-I-interferon signaling drives microglial dysfunction and senescence in human iPSC models of Down syndrome and Alzheimer’s disease. Cell Stem Cell 29:1135–1153.e8. doi: 10.1016/j.stem.2022.06.007

28. Kanehisa M, Furumichi M, Sato Y, Ishiguro-Watanabe M, Tanabe M (2021) KEGG: integrating viruses and cellular organisms. Nucleic Acids Res 49:D545–D551. doi: 10.1093/nar/gkaa970

29. Keren-Shaul H, Spinrad A, Weiner A, Matcovitch-Natan O, Dvir-Szternfeld R, Ulland TK, David E, Baruch K, Lara-Astaiso D, Toth B, Itzkovitz S, Colonna M, Schwartz M, Amit I (2017) A Unique Microglia Type Associated with Restricting Development of Alzheimer’s Disease. Cell 169:1276–1290.e17. doi: 10.1016/j.cell.2017.05.018

30. Kraft AW, Hu X, Yoon H, Yan P, Xiao Q, Wang Y, Gil SC, Brown J, Wilhelmsson U, Restivo JL, Cirrito JR, Holtzman DM, Kim J, Pekny M, Lee J-M (2013) Attenuating astrocyte activation accelerates plaque pathogenesis in APP/PS1 mice. FASEB J 27:187–98. doi: 10.1096/fj.12-208660

31. Kunkle BW, Grenier-Boley B, Sims R, Bis JC, Damotte V, Naj AC, Boland A, Vronskaya M, van der Lee SJ, Amlie-Wolf A, Bellenguez C, Frizatti A, Chouraki V, Martin ER, Sleegers K, Badarinarayan N, Jakobsdottir J, Hamilton-Nelson KL, Moreno-Grau S, Olaso R, Raybould R, Chen Y, Kuzma AB, Hiltunen M, Morgan T, Ahmad S, Vardarajan BN, Epelbaum J, Hoffmann P, Boada M, Beecham GW, Garnier J-G, Harold D, Fitzpatrick AL, Valladares O, Moutet M-L, Gerrish A, Smith A V., Qu L, Bacq D, Denning N, Jian X, Zhao Y, Del Zompo M, Fox NC, Choi S-H, Mateo I, Hughes JT, Adams HH, Malamon J, Sanchez-Garcia F, Patel Y, Brody JA, Dombroski BA, Naranjo MCD, Daniilidou M, Eiriksdottir G, Mukherjee S, Wallon D, Uphill J, Aspelund T, Cantwell LB, Garzia F, Galimberti D, Hofer E, Butkiewicz M, Fin B, Scarpini E, Sarnowski C, Bush WS, Meslage S, Kornhuber J, White CC, Song Y, Barber RC, Engelborghs S, Sordon S, Voijnovic D, Adams PM, Vandenberghe R, Mayhaus M, Cupples LA, Albert MS, De Deyn PP, Gu W, Himali JJ, Beekly D, Squassina A, Hartmann AM, Orellana A, Blacker D, Rodriguez-Rodriguez E, Lovestone S, Garcia ME, Doody RS, Munoz-Fernadez C, Sussams R, Lin H, Fairchild TJ, Benito YA, Holmes C, Karamujić-Čomić H, Frosch MP, Thonberg H, Maier W, Roshchupkin G, Ghetti B, Giedraitis V, Kawalia A, Li S, Huebinger RM, Kilander L, Moebus S, Hernández I, Kamboh MI, Brundin R, Turton J, Yang Q, Katz MJ, Concari L, Lord J, Beiser AS, Keene CD, Helisalmi S, Kloszewska I, Kukull WA, Koivisto AM, Lynch A, Tarraga L, Larson EB, Haapasalo A, Lawlor B, Mosley TH, Lipton RB, Solfrizzi V, Gill M, Longstreth WT, Montine TJ, Frisardi V, Diez-Fairen M, Rivadeneira F, Petersen RC, Deramecourt V, Alvarez I, Salani F, Ciaramella A, Boerwinkle E, Reiman EM, Fievet N, Rotter JI, Reisch JS, Hanon O, Cupidi C, Andre Uitterlinden AG, Royall DR, Dufouil C, Maletta RG, de Rojas I, Sano M, Brice A, Cecchetti R, George-Hyslop PS, Ritchie K, Tsolaki M, Tsuang DW, Dubois B, Craig D, Wu C-K, Soininen H, Avramidou D, Albin RL, Fratiglioni L, Germanou A, Apostolova LG, Keller L, Koutroumani M, Arnold SE, Panza F, Gkatzima O, Asthana S, Hannequin D, Whitehead P, Atwood CS, Caffarra P, Hampel H, Quintela I, Carracedo Á, Lannfelt L, Rubinsztein DC, Barnes LL, Pasquier F, Frölich L, Barral S, McGuinness B, Beach TG, Johnston JA, Becker JT, Passmore P, Bigio EH, Schott JM, Bird TD, Warren JD, Boeve BF, Lupton MK, Bowen JD, Proitsi P, Boxer A, Powell JF, Burke JR, Kauwe JSK, Burns JM, Mancuso M, Buxbaum JD, Bonuccelli U, Cairns NJ, McQuillin A, Cao C, Livingston G, Carlson CS, Bass NJ, Carlsson CM, Hardy J, Carney RM, Bras J, Carrasquillo MM, Guerreiro R, Allen M, Chui HC, Fisher E, Masullo C, Crocco EA, DeCarli C, Bisceglio G, Dick M, Ma L, Duara R, Graff-Radford NR, Evans DA, Hodges A, Faber KM, Scherer M, Fallon KB, Riemenschneider M, Fardo DW, Heun R, Farlow MR, Kölsch H, Ferris S, Leber M, Foroud TM, Heuser I, Galasko DR, Giegling I, Gearing M, Hüll M, Geschwind DH, Gilbert JR, Morris J, Green RC, Mayo K, Growdon JH, Feulner T, Hamilton RL, Harrell LE, Drichel D, Honig LS, Cushion TD, Huentelman MJ, Hollingworth P, Hulette CM, Hyman BT, Marshall R, Jarvik GP, Meggy A, Abner E, Menzies GE, Jin L-W, Leonenko G, Real LM, Jun GR, Baldwin CT, Grozeva D, Karydas A, Russo G, Kaye JA, Kim R, Jessen F, Kowall NW, Vellas B, Kramer JH, Vardy E, LaFerla FM, Jöckel K-H, Lah JJ, Dichgans M, Leverenz JB, Mann D, Levey AI, Pickering-Brown S, Lieberman AP, Klopp N, Lunetta KL, Wichmann H-E, Lyketsos CG, Morgan K, Marson DC, Brown K, Martiniuk F, Medway C, Mash DC, Nöthen MM, Masliah E, Hooper NM, McCormick WC, Daniele A, McCurry SM, Bayer A, McDavid AN, Gallacher J, McKee AC, van den Bussche H, Mesulam M, Brayne C, Miller BL, Riedel-Heller S, Miller CA, Miller JW, Al-Chalabi A, Morris JC, Shaw CE, Myers AJ, Wiltfang J, O’Bryant S, Olichney JM, Alvarez V, Parisi JE, Singleton AB, Paulson HL, Collinge J, Perry WR, Mead S, Peskind E, Cribbs DH, Rossor M, Pierce A, Ryan NS, Poon WW, Nacmias B, Potter H, Sorbi S, Quinn JF, Sacchinelli E, Raj A, Spalletta G, Raskind M, Caltagirone C, Bossù P, Orfei MD, Reisberg B, Clarke R, Reitz C, Smith AD, Ringman JM, Warden D, Roberson ED, Wilcock G, Rogaeva E, Bruni AC, Rosen HJ, Gallo M, Rosenberg RN, Ben-Shlomo Y, Sager MA, Mecocci P, Saykin AJ, Pastor P, Cuccaro ML, Vance JM, Schneider JA, Schneider LS, Slifer S, Seeley WW, Smith AG, Sonnen JA, Spina S, Stern RA, Swerdlow RH, Tang M, Tanzi RE, Trojanowski JQ, Troncoso JC, Van Deerlin VM, Van Eldik LJ, Vinters H V., Vonsattel JP, Weintraub S, Welsh-Bohmer KA, Wilhelmsen KC, Williamson J, Wingo TS, Woltjer RL, Wright CB, Yu C-E, Yu L, Saba Y, Pilotto A, Bullido MJ, Peters O, Crane PK, Bennett D, Bosco P, Coto E, Boccardi V, De Jager PL, Lleo A, Warner N, Lopez OL, Ingelsson M, Deloukas P, Cruchaga C, Graff C, Gwilliam R, Fornage M, Goate AM, Sanchez-Juan P, Kehoe PG, Amin N, Ertekin-Taner N, Berr C, Debette S, Love S, Launer LJ, Younkin SG, Dartigues J-F, Corcoran C, Ikram MA, Dickson DW, Nicolas G, Campion D, Tschanz J, Schmidt H, Hakonarson H, Clarimon J, Munger R, Schmidt R, Farrer LA, Van Broeckhoven C, C. O’Donovan M, DeStefano AL, Jones L, Haines JL, Deleuze J-F, Owen MJ, Gudnason V, Mayeux R, Escott-Price V, Psaty BM, Ramirez A, Wang L-S, Ruiz A, van Duijn CM, Holmans PA, Seshadri S, Williams J, Amouyel P, Schellenberg GD, Lambert J-C, Pericak-Vance MA (2019) Genetic meta-analysis of diagnosed Alzheimer’s disease identifies new risk loci and implicates Aβ, tau, immunity and lipid processing. Nat Genet 51:414–430. doi: 10.1038/s41588-019-0358-2

32. Lee H, Pearse R V, Lish AM, Pan C, Augur ZM, Terzioglu G, Gaur P, Liao M, Fujita M, Tio ES, Duong DM, Felsky D, Seyfried NT, Menon V, Bennett DA, De Jager PL, Young-Pearse TL (2024) Contributions of genetic variation in astrocytes to cell and molecular mechanisms of risk and resilience to late onset Alzheimer’s disease. bioRxiv. doi: 10.1101/2024.07.31.605928

33. Lee S-H, Meilandt WJ, Xie L, Gandham VD, Ngu H, Barck KH, Rezzonico MG, Imperio J, Lalehzadeh G, Huntley MA, Stark KL, Foreman O, Carano RAD, Friedman BA, Sheng M, Easton A, Bohlen CJ, Hansen D V (2021) Trem2 restrains the enhancement of tau accumulation and neurodegeneration by β-amyloid pathology. Neuron 109:1283–1301.e6. doi: 10.1016/j.neuron.2021.02.010

34. Lefebvre-Omar C, Liu E, Dalle C, D’Incamps BL, Bigou S, Daube C, Karpf L, Davenne M, Robil N, Jost Mousseau C, Blanchard S, Tournaire G, Nicaise C, Salachas F, Lacomblez L, Seilhean D, Lobsiger CS, Millecamps S, Boillée S, Bohl D (2023) Neurofilament accumulations in amyotrophic lateral sclerosis patients’ motor neurons impair axonal initial segment integrity. Cell Mol Life Sci 80:150. doi: 10.1007/s00018-023-04797-6

35. Levy-Lahad E, Wasco W, Poorkaj P, Romano D, Oshima J, Pettingell W, Yu C, Jondro P, Schmidt S, Wang K, Al. E (1995) Candidate gene for the chromosome 1 familial Alzheimer’s disease locus. Science (1979) 269:973–977. doi: 10.1126/science.7638622

36. Leyns CEG, Gratuze M, Narasimhan S, Jain N, Koscal LJ, Jiang H, Manis M, Colonna M, Lee VMY, Ulrich JD, Holtzman DM (2019) TREM2 function impedes tau seeding in neuritic plaques. Nat Neurosci 22:1217–1222. doi: 10.1038/s41593-019-0433-0

37. Liddelow SA, Guttenplan KA, Clarke LE, Bennett FC, Bohlen CJ, Schirmer L, Bennett ML, Münch AE, Chung WS, Peterson TC, Wilton DK, Frouin A, Napier BA, Panicker N, Kumar M, Buckwalter MS, Rowitch DH, Dawson VL, Dawson TM, Stevens B, Barres BA (2017) Neurotoxic reactive astrocytes are induced by activated microglia. Nature 541:481–487. doi: 10.1038/nature21029

38. Londino JD, Gulick D, Isenberg JS, Mallampalli RK (2015) Cleavage of Signal Regulatory Protein α (SIRPα) Enhances Inflammatory Signaling. Journal of Biological Chemistry 290:31113–31125. doi: 10.1074/jbc.M115.682914

39. Love MI, Huber W, Anders S (2014) Moderated estimation of fold change and dispersion for RNA-seq data with DESeq2. Genome Biol 15:550. doi: 10.1186/s13059-014-0550-8

40. Luo W, Brouwer C (2013) Pathview: an R/Bioconductor package for pathway-based data integration and visualization. Bioinformatics 29:1830–1831. doi: 10.1093/bioinformatics/btt285

41. Mathys H, Boix CA, Akay LA, Xia Z, Davila-Velderrain J, Ng AP, Jiang X, Abdelhady G, Galani K, Mantero J, Band N, James BT, Babu S, Galiana-Melendez F, Louderback K, Prokopenko D, Tanzi RE, Bennett DA, Tsai L-H, Kellis M (2024) Single-cell multiregion dissection of Alzheimer’s disease. Nature 632:858–868. doi: 10.1038/s41586-024-07606-7

42. Mawuenyega KG, Sigurdson W, Ovod V, Munsell L, Kasten T, Morris JC, Yarasheski KE, Bateman RJ (2010) Decreased Clearance of CNS β-Amyloid in Alzheimer’s Disease. Science (1979) 330:1774–1774. doi: 10.1126/science.1197623

43. Merilahti JAM, Elenius K (2019) Gamma-secretase-dependent signaling of receptor tyrosine kinases. Oncogene 38:151–163. doi: 10.1038/s41388-018-0465-z

44. Montoliu-Gaya L, Alcolea D, Ashton NJ, Pegueroles J, Levin J, Bosch B, Lantero-Rodriguez J, Carmona-Iragui M, Wagemann O, Balasa M, Kac PR, Barroeta I, Lladó A, Brum WS, Videla L, Gonzalez-Ortiz F, Benejam B, Arranz Martínez JJ, Karikari TK, Nübling G, Bejanin A, Benedet AL, Blesa R, Lleó A, Blennow K, Sánchez-Valle R, Zetterberg H, Fortea J (2023) Plasma and cerebrospinal fluid glial fibrillary acidic protein levels in adults with Down syndrome: a longitudinal cohort study. EBioMedicine 90:104547. doi: 10.1016/j.ebiom.2023.104547

45. Oksanen M, Petersen AJ, Naumenko N, Puttonen K, Lehtonen Š, Gubert Olivé M, Shakirzyanova A, Leskelä S, Sarajärvi T, Viitanen M, Rinne JO, Hiltunen M, Haapasalo A, Giniatullin R, Tavi P, Zhang SC, Kanninen KM, Hämäläinen RH, Koistinaho J (2017) PSEN1 Mutant iPSC-Derived Model Reveals Severe Astrocyte Pathology in Alzheimer’s Disease. Stem Cell Reports. doi: 10.1016/j.stemcr.2017.10.016

46. Pink AE, Simpson MA, Desai N, Trembath RC, Barker JNW (2013) γ-Secretase mutations in hidradenitis suppurativa: new insights into disease pathogenesis. J Invest Dermatol 133:601–607. doi: 10.1038/jid.2012.372

47. Portelius E, Appelkvist P, Strömberg K, Höglund K (2014) Characterization of the effect of a novel γ-secretase modulator on Aβ: A clinically translatable model. Curr Pharm Des 20:2484–2490

48. Potier M-C, Steiner H, Chávez-Gutiérrez L (2024) Amyloid β, γ-secretase, and familial Alzheimer’s disease. Lancet Neurol 23:852–853. doi: 10.1016/S1474-4422(24)00292-8

49. Potter R, Patterson BW, Elbert DL, Ovod V, Kasten T, Sigurdson W, Mawuenyega K, Blazey T, Goate A, Chott R, Yarasheski KE, Holtzman DM, Morris JC, Benzinger TLS, Bateman RJ (2013) Increased in vivo amyloid-b42 production, exchange, and loss in presenilin mutation carriers. Sci Transl Med 5. doi: 10.1126/scitranslmed.3005615

50. Prakash P, Erdjument-Bromage H, O’Dea MR, Munson CN, Labib D, Fossati V, Neubert TA, Liddelow SA (2023) Proteomic profiling of interferon-responsive reactive astrocytes in rodent and human. Glia 625–642. doi: 10.1002/glia.24494

51. Ryan NS, Nicholas JM, Weston PSJ, Liang Y, Lashley T, Guerreiro R, Adamson G, Kenny J, Beck J, Chavez-Gutierrez L, de Strooper B, Revesz T, Holton J, Mead S, Rossor MN, Fox NC (2016) Clinical phenotype and genetic associations in autosomal dominant familial Alzheimer’s disease: a case series. Lancet Neurol 15:1326–1335. doi: 10.1016/S1474-4422(16)30193-4

52. Ryman DC, Acosta-Baena N, Aisen PS, Bird T, Danek A, Fox NC, Goate A, Frommelt P, Ghetti B, Langbaum JBS, Lopera F, Martins R, Masters CL, Mayeux RP, McDade E, Moreno S, Reiman EM, Ringman JM, Salloway S, Schofield PR, Sperling R, Tariot PN, Xiong C, Morris JC, Bateman RJ (2014) Symptom onset in autosomal dominant Alzheimer disease: A systematic review and meta-analysis. Neurology 83:253–260. doi: 10.1212/WNL.0000000000000596

53. Salih DA, Bayram S, Guelfi S, Reynolds RH, Shoai M, Ryten M, Brenton JW, Zhang D, Matarin M, Botia JA, Shah R, Brookes KJ, Guetta-Baranes T, Morgan K, Bellou E, Cummings DM, Escott-Price V, Hardy J (2019) Genetic variability in response to amyloid beta deposition influences Alzheimer’s disease risk. Brain Commun 1. doi: 10.1093/braincomms/fcz022

54. Scheinin I, Sie D, Bengtsson H, Van De Wiel MA, Olshen AB, Van Thuijl HF, Van Essen HF, Eijk PP, Rustenburg F, Meijer GA, Reijneveld JC, Wesseling P, Pinkel D, Albertson DG, Ylstra B (2014) DNA copy number analysis of fresh and formalin-fixed specimens by shallow whole-genome sequencing with identification and exclusion of problematic regions in the genome assembly. Genome Res 24:2022–2032. doi: 10.1101/gr.175141.114

55. Schubert M, Klinger B, Klünemann M, Sieber A, Uhlitz F, Sauer S, Garnett MJ, Blüthgen N, Saez-Rodriguez J (2018) Perturbation-response genes reveal signaling footprints in cancer gene expression. Nat Commun 9:20. doi: 10.1038/s41467-017-02391-6

56. Schultz SA, Liu L, Schultz AP, Fitzpatrick CD, Levin R, Bellier J-P, Shirzadi Z, Joseph-Mathurin N, Chen CD, Benzinger TLS, Day GS, Farlow MR, Gordon BA, Hassenstab JJ, Jack CR, Jucker M, Karch CM, Lee J-H, Levin J, Perrin RJ, Schofield PR, Xiong C, Johnson KA, McDade E, Bateman RJ, Sperling RA, Selkoe DJ, Chhatwal JP, Dominantly Inherited Alzheimer Network (2024) γ-Secretase activity, clinical features, and biomarkers of autosomal dominant Alzheimer’s disease: cross-sectional and longitudinal analysis of the Dominantly Inherited Alzheimer Network observational study (DIAN-OBS). Lancet Neurol 23:913–924. doi: 10.1016/S1474-4422(24)00236-9

57. Serrano-Pozo A, Muzikansky A, Gómez-Isla T, Growdon JH, Betensky RA, Frosch MP, Hyman BT (2013) Differential relationships of reactive astrocytes and microglia to fibrillar amyloid deposits in alzheimer disease. J Neuropathol Exp Neurol 72:462– 471. doi: 10.1097/NEN.0b013e3182933788

58. Setó-Salvia N, Esteras N, de Silva R, de Pablo-Fernandez E, Arber C, Toomey CE, Polke JM, Morris HR, Rohrer JD, Abramov AY, Patani R, Wray S, Warner TT (2022) Elevated 4R-tau in astrocytes from asymptomatic carriers of the MAPT 10+16 intronic mutation. J Cell Mol Med 26:1327–1331. doi: 10.1111/jcmm.17136

59. Shen Y, Ali M, Timsina J, Wang C, Do A, Western D, Liu M, Gorijala P, Budde J, Liu H, Gordon B, Mcdade E, Morris JC, Llibre-Guerra JJ, Bateman RJ, Joseph-Mathurin N, Perrin RJ, Maschi D, Wyss-Coray T, Pastor P, Goate A, Renton AE, Surace EI, Johnson ECB, Levey AI, Alvarez I, Levin J, Ringman JM, Allegri RF, Seyfried N, Day GS, Wu Q, Fernández MV, Network A, Ibanez L, Sung YJ, Cruchaga C Systematic proteomics in Autosomal dominant Alzheimer’s disease reveals decades-early changes of CSF proteins in neuronal death, and immune pathways. MedrXiv. doi: 10.1101/2024.01.12.24301242

60. Sherrington R, Rogaev EI, Liang Y, Rogaeva EA, Levesque G, Ikeda M, Chi H, Lin C, Li G, Holman K, Tsuda T, Mar L, Foncin J-F, Bruni AC, Montesi MP, Sorbi S, Rainero I, Pinessi L, Nee L, Chumakov I, Pollen D, Brookes A, Sanseau P, Polinsky RJ, Wasco W, Da Silva HAR, Haines JL, Pericak-Vance MA, Tanzi RE, Roses AD, Fraser PE, Rommens JM, St George-Hyslop PH (1995) Cloning of a gene bearing missense mutations in early-onset familial Alzheimer’s disease. Nature 375:754–760. doi: 10.1038/375754a0

61. Siman R, Reaume AG, Savage MJ, Trusko S, Lin YG, Scott RW, Flood DG (2000) Presenilin-1 P264L knock-in mutation: Differential effects on Aβ production, amyloid deposition, and neuronal vulnerability. Journal of Neuroscience 20:8717–8726. doi: 10.1523/jneurosci.20-23-08717.2000

62. Struhl G, Adachi A (2000) Requirements for Presenilin-Dependent Cleavage of Notch and Other Transmembrane Proteins. Mol Cell 6:625–636. doi: 10.1016/S1097-2765(00)00061-7

63. Szaruga M, Munteanu B, Lismont S, Veugelen S, Horré K, Mercken M, Saido TC, Ryan NS, De Vos T, Savvides SN, Gallardo R, Schymkowitz J, Rousseau F, Fox NC, Hopf C, De Strooper B, Chávez-Gutiérrez L (2017) Alzheimer’s-Causing Mutations Shift Aβ Length by Destabilizing γ-Secretase-Aβn Interactions. Cell 170:443–456.e14. doi: 10.1016/j.cell.2017.07.004

64. Tyzack G, Lakatos A, Patani R (2016) Human Stem Cell-Derived Astrocytes: Specification and Relevance for Neurological Disorders. Curr Stem Cell Rep 2:236– 247. doi: 10.1007/s40778-016-0049-1

65. Wang B, Yang W, Wen W, Sun J, Su B, Liu B, Ma D, Lv D, Wen Y, Qu T, Chen M, Sun M, Shen Y, Zhang X (2010) Γ-Secretase Gene Mutations in Familial Acne Inversa. Science (1979) 330:1065. doi: 10.1126/science.1196284

66. Willumsen N, Arber C, Lovejoy C, Toombs J, Alatza A, Weston PSJ, Chávez-Gutiérrez L, Hardy J, Zetterberg H, Fox NC, Ryan NS, Lashley T, Wray S (2022) The PSEN1 E280G mutation leads to increased amyloid-β43 production in induced pluripotent stem cell neurons and deposition in brain tissue. Brain Commun 5. doi: 10.1093/braincomms/fcac321

67. Yang HS, Teng L, Kang D, Menon V, Ge T, Finucane HK, Schultz AP, Properzi M, Klein HU, Chibnik LB, Schneider JA, Bennett DA, Hohman TJ, Mayeux RP, Johnson KA, De Jager PL, Sperling RA (2023) Cell-type-specific Alzheimer’s disease polygenic risk scores are associated with distinct disease processes in Alzheimer’s disease. Nat Commun 14:1–13. doi: 10.1038/s41467-023-43132-2

68. Zurhove K, Nakajima C, Herz J, Bock HH, May P (2008) γ-Secretase Limits the Inflammatory Response Through the Processing of LRP1. Sci Signal 1. doi: 10.1126/scisignal.1164263

