## Supplementary figures for "Mutations in *PSEN1* predispose inflammation in an astrocyte model of familial Alzheimer’s disease through disrupted regulated intramembrane proteolysis"

### Slide 1
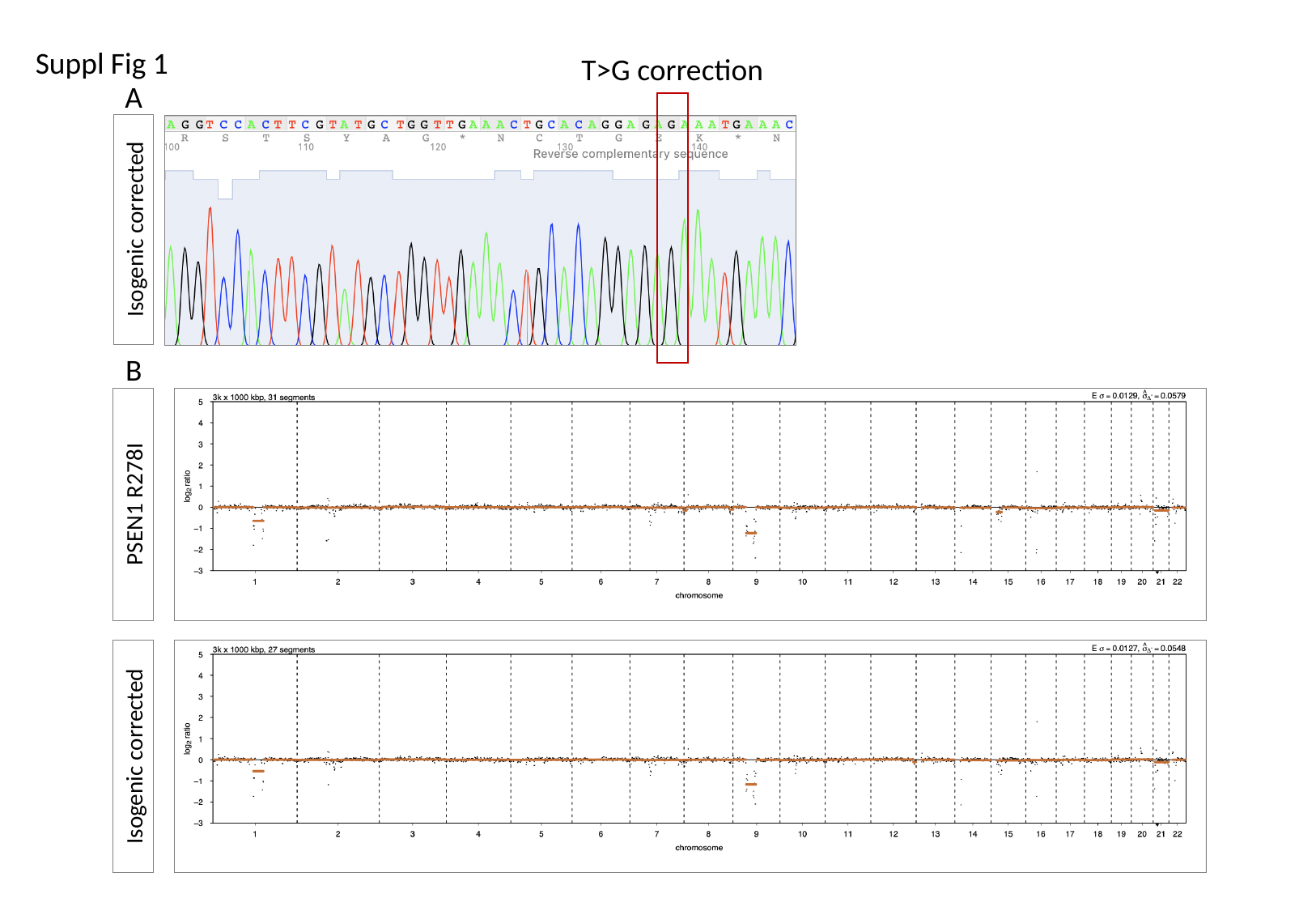

Suppl Fig 1
T>G correction
A
Isogenic corrected
B
PSEN1 R278I
Isogenic corrected

### Slide 2
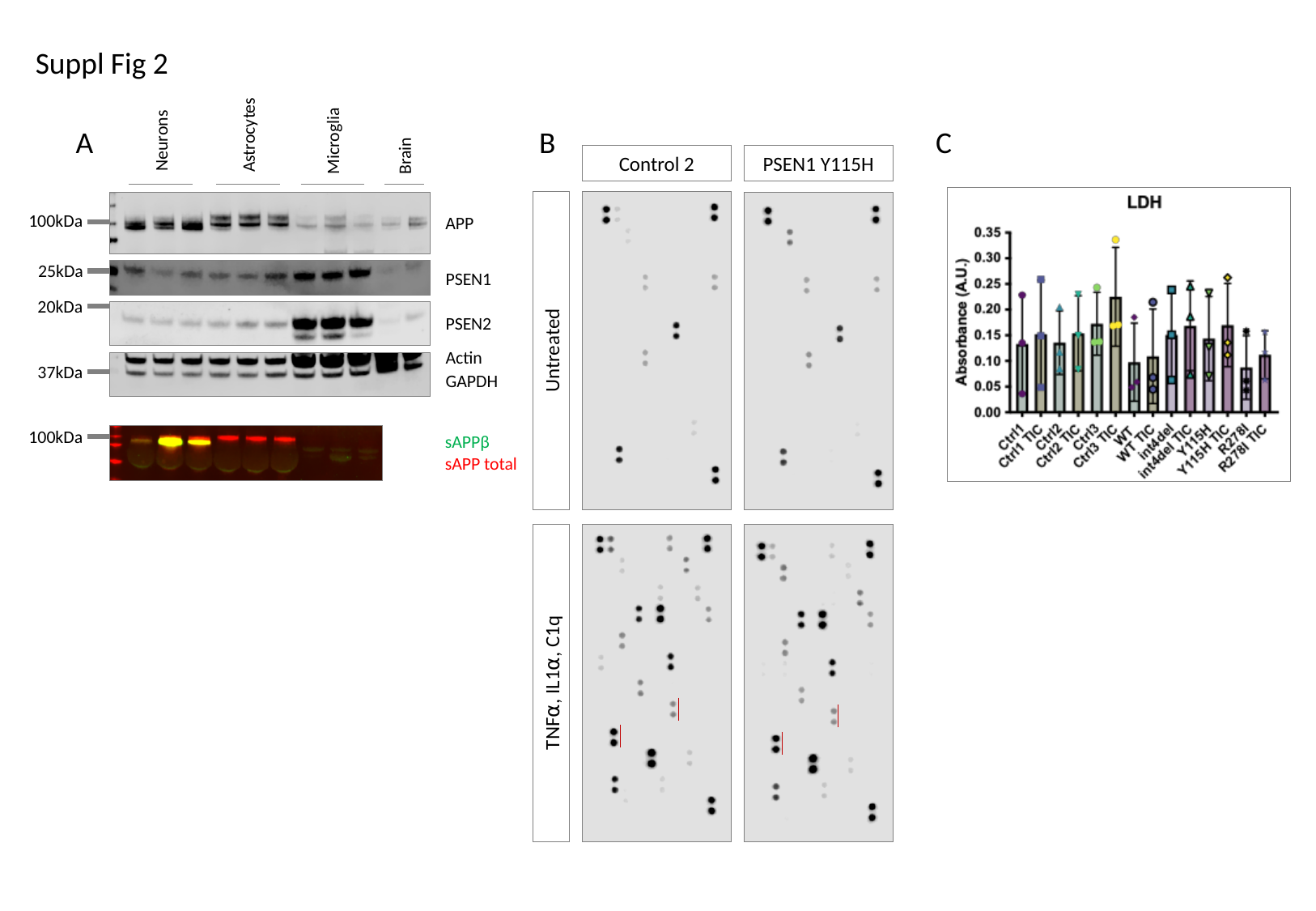

Suppl Fig 2
A
B
C
Astrocytes
Neurons
Microglia
Brain
Control 2
PSEN1 Y115H
100kDa
APP
25kDa
PSEN1
20kDa
PSEN2
Untreated
Actin
37kDa
GAPDH
100kDa
sAPPβ
sAPP total
TNF⍺, IL1⍺, C1q

### Slide 3
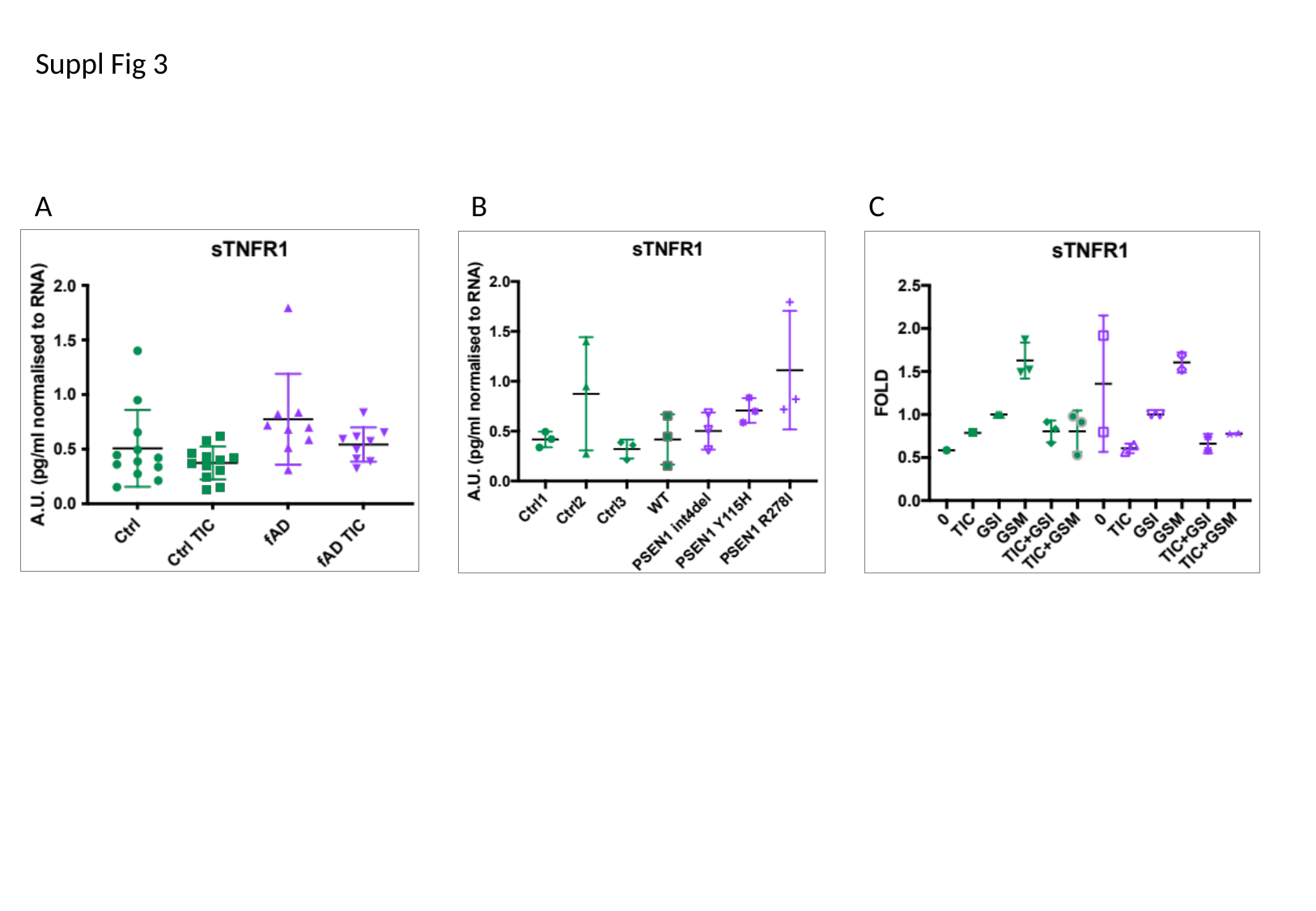

Suppl Fig 3
A
B
C

### Slide 4
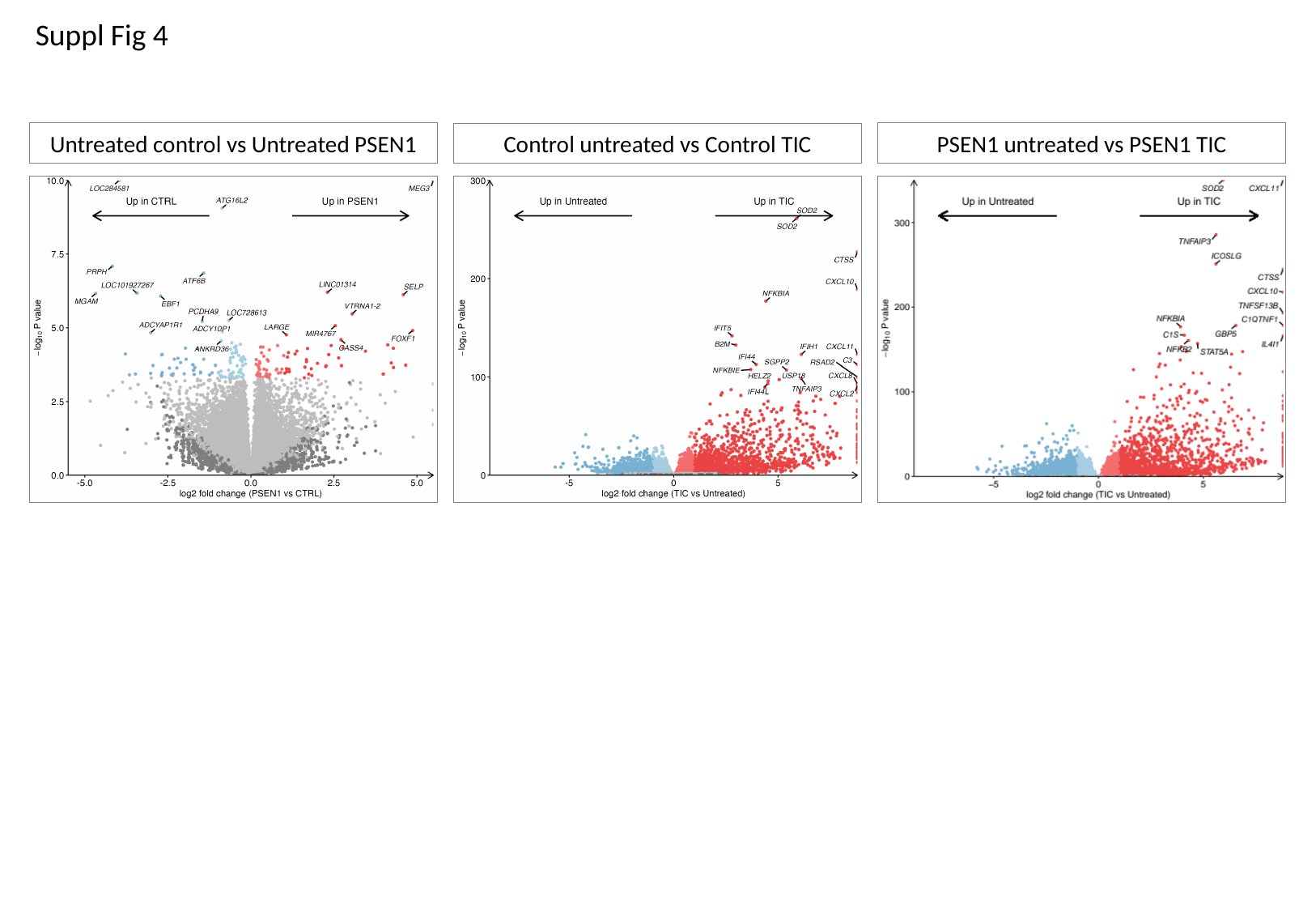

Suppl Fig 4
Untreated control vs Untreated PSEN1
PSEN1 untreated vs PSEN1 TIC
Control untreated vs Control TIC

### Slide 5
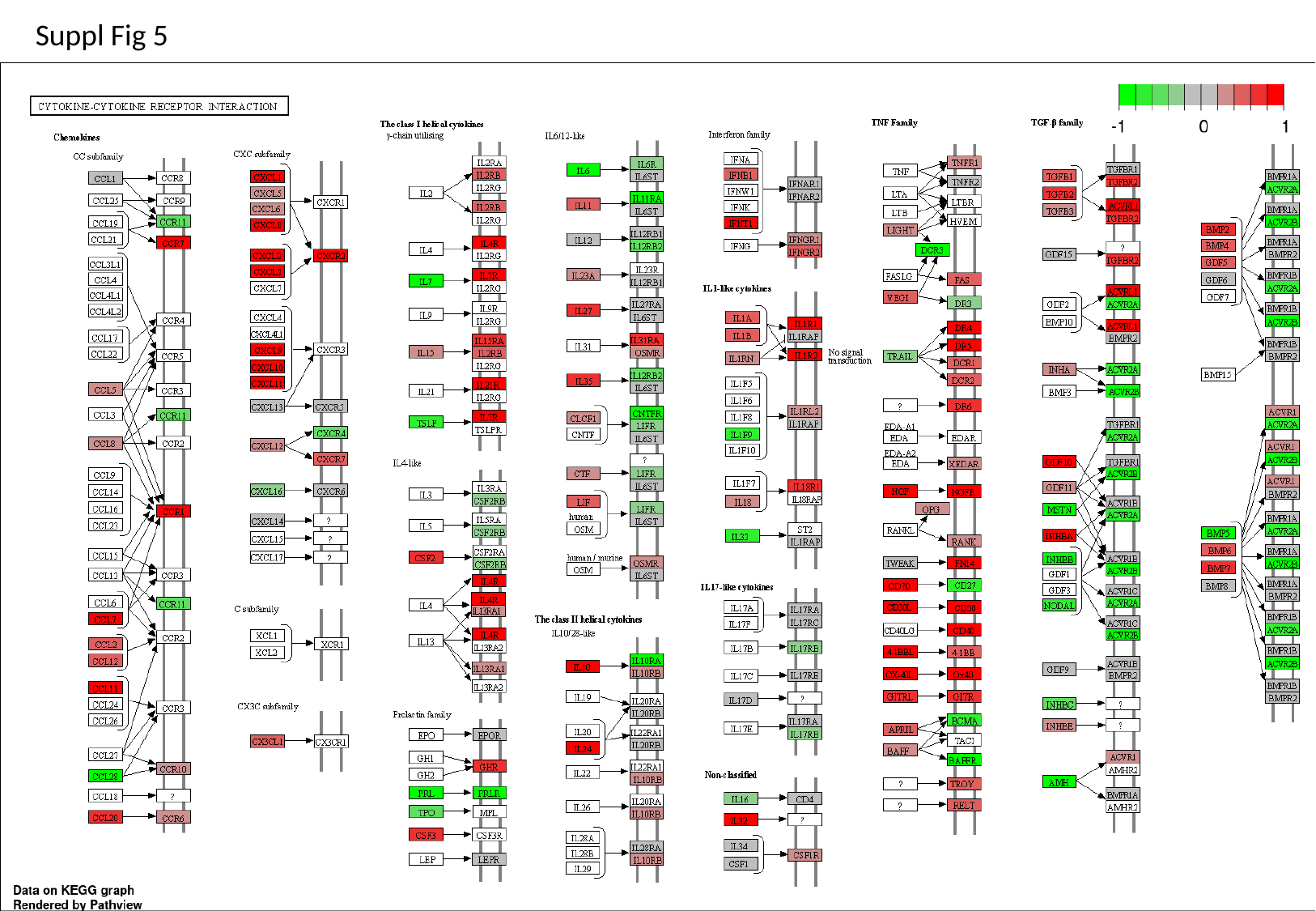

Suppl Fig 5

### Slide 6
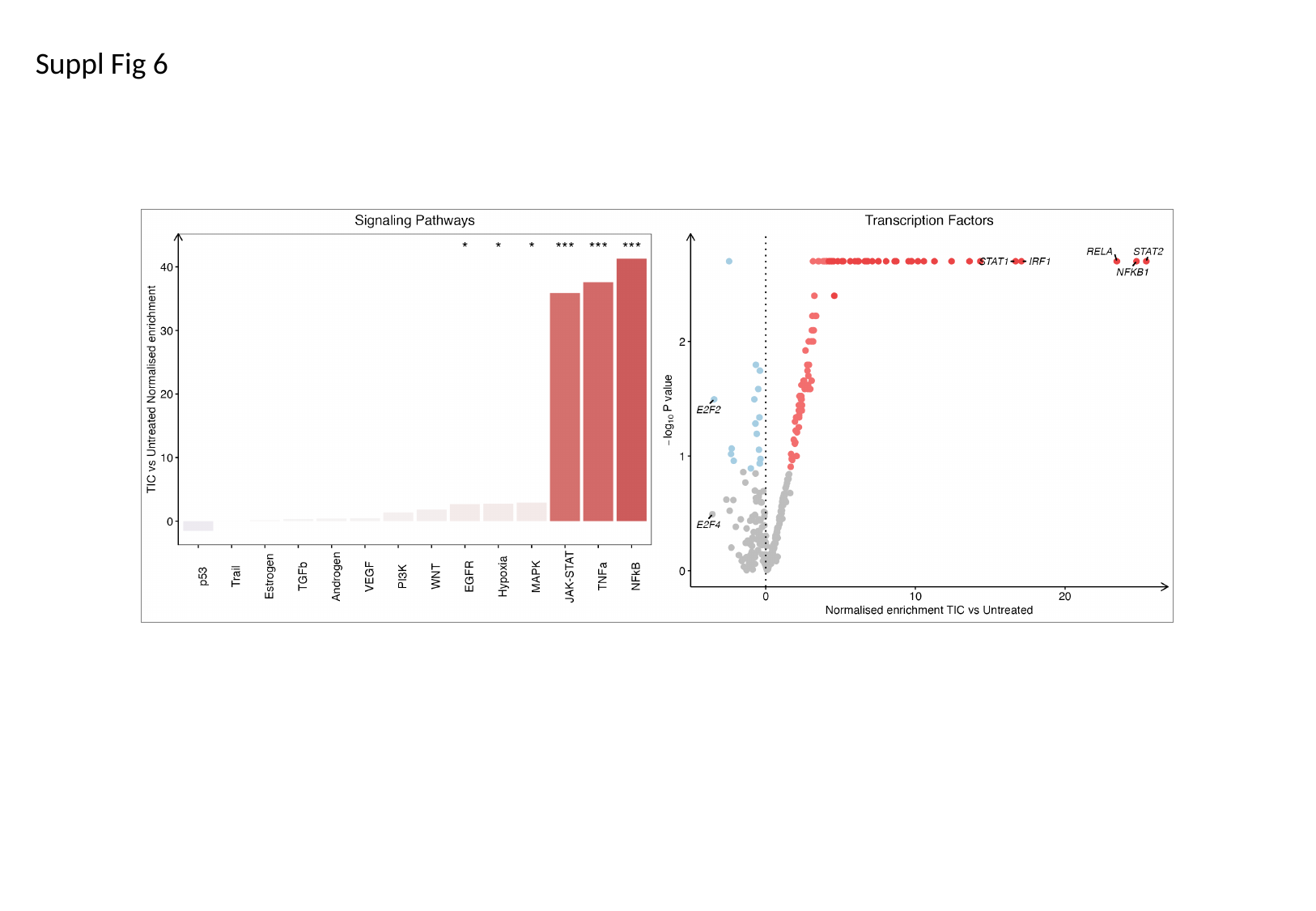

Suppl Fig 6

### Slide 7
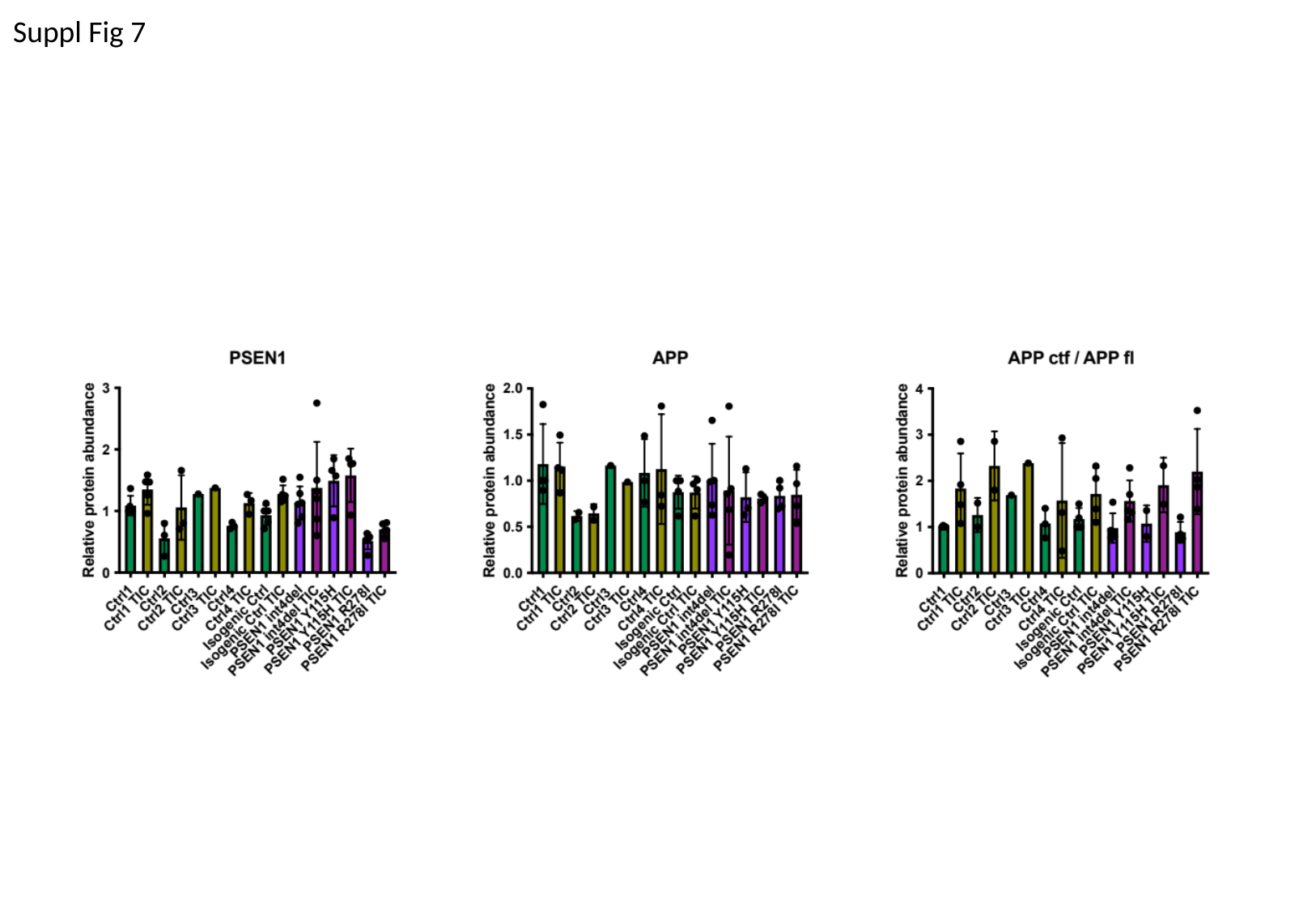

Suppl Fig 7

### Slide 8
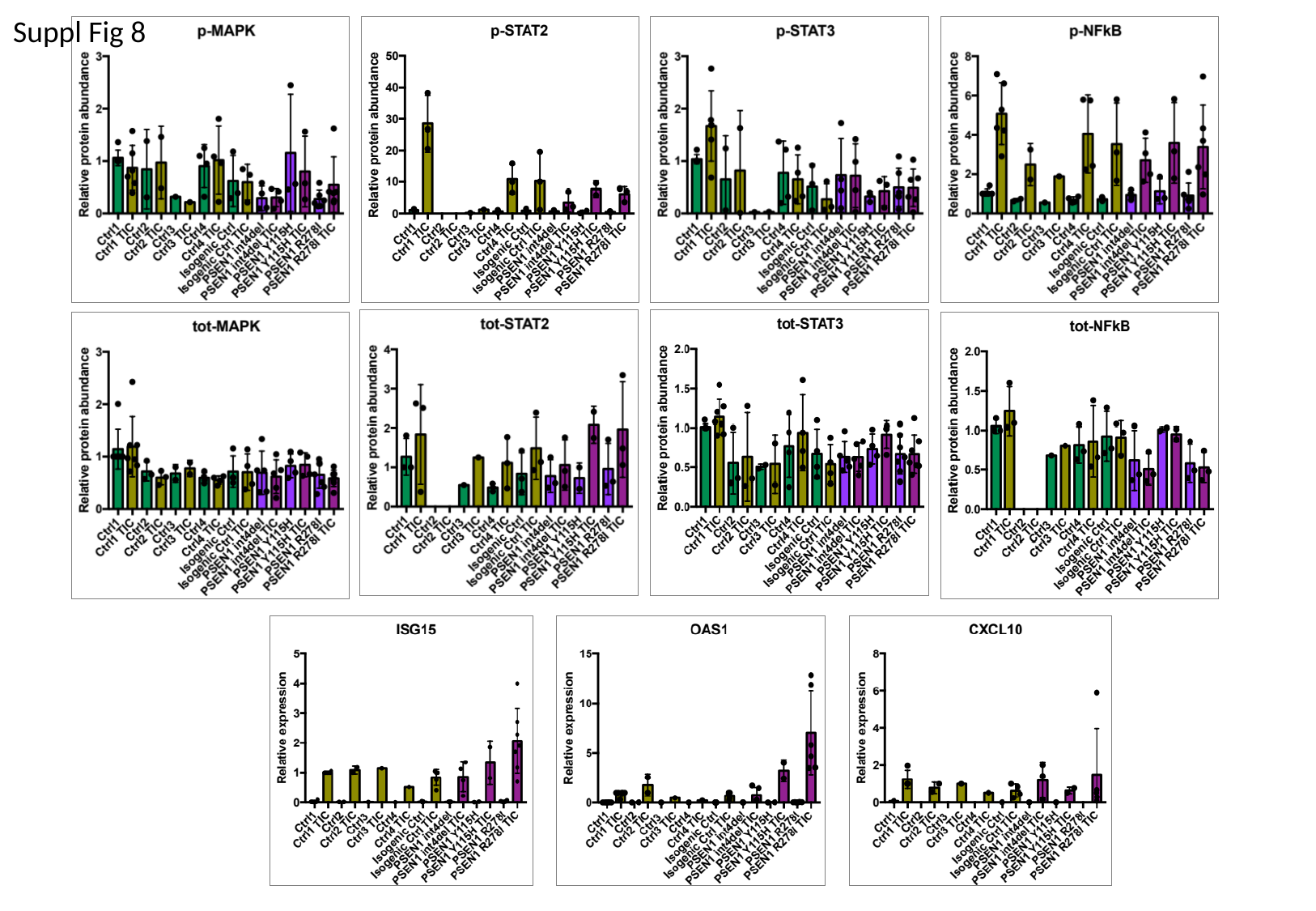

Suppl Fig 8
